# M-Phase oscillations in PP1 activity are essential for accurate progression through mammalian oocyte meiosis

**DOI:** 10.1101/2021.09.28.461976

**Authors:** Nicole J. Camlin, Ilakkiya Venkatachalam, Janice P. Evans

**Affiliations:** Department of Biological Sciences, Purdue University, West Lafayette, IN 47907

**Keywords:** Oocyte, meiosis, PP1, M-Phase

## Abstract

Tightly controlled fluctuations in kinase and phosphatase activity play important roles in regulating M-Phase transitions. Protein Phosphatase 1 (PP1) is one of these phosphatases, with oscillations in PP1 activity driving mitotic M-Phase. Evidence from a variety of experimental systems also points to roles in meiosis. Here we report that PP1 is important for M-Phase transitions through mouse oocyte meiosis. We employed a unique small-molecule approach to inhibit or activate PP1 at distinct phases of mouse oocyte meiosis. These studies show that temporal control of PP1 activity is essential for G2/M transition, metaphase I/anaphase I transition, and the formation of a normal metaphase II oocyte. Our data also reveal that inappropriate activation of PP1 is more deleterious at G2/M transition than at prometaphase I-to-metaphase I, and that an active pool of PP1 during prometaphase is vital for metaphase I/anaphase I transition and metaphase II chromosome alignment. Taken together, these results establish that loss of oscillations in PP1 activity causes a range of severe meiotic defects, pointing to essential roles for PP1 in female fertility, and more broadly, M-Phase regulation.

**Summary statement:** Altering the normal cyclical activity of the phosphatase PP1 in oocytes causes a range of severe meiotic defects, pointing to essential roles for PP1 in M-Phase entry, progression, and exit.

## Introduction

Accurate oocyte meiosis is essential for female fertility and the development of healthy offspring. Paradoxically, mammalian oocyte meiosis is inherently error-prone, with aneuploidy observed in 10-30% of human oocytes, making aneuploidy a leading cause of miscarriage and birth defects (Hassold and Hunt, 2001; Nagaoka et al., 2012). Key to successful meiosis is precise M-Phase regulation driven by widespread changes in protein phosphorylation (Fig. 1). This dynamic phospho-environment is driven by tightly controlled fluctuations in both kinase and phosphatase activity. CDK1 is largely responsible for the increase in protein phosphorylation that propels M-Phase entry and progression (Adhikari et al., 2012; Crncec and Hochegger, 2019). Phosphatases, including the closely related serine/threonine phosphatases Protein Phosphatase 1 (PP1) and Protein Phosphatase 2A (PP2A), regulate M-Phase exit by removing phosphorylations needed to maintain M-Phase and act as the major phosphatases that oppose CDK1 (St-Denis et al., 2016) (Fig. 1A-B).

**Figure 1:**
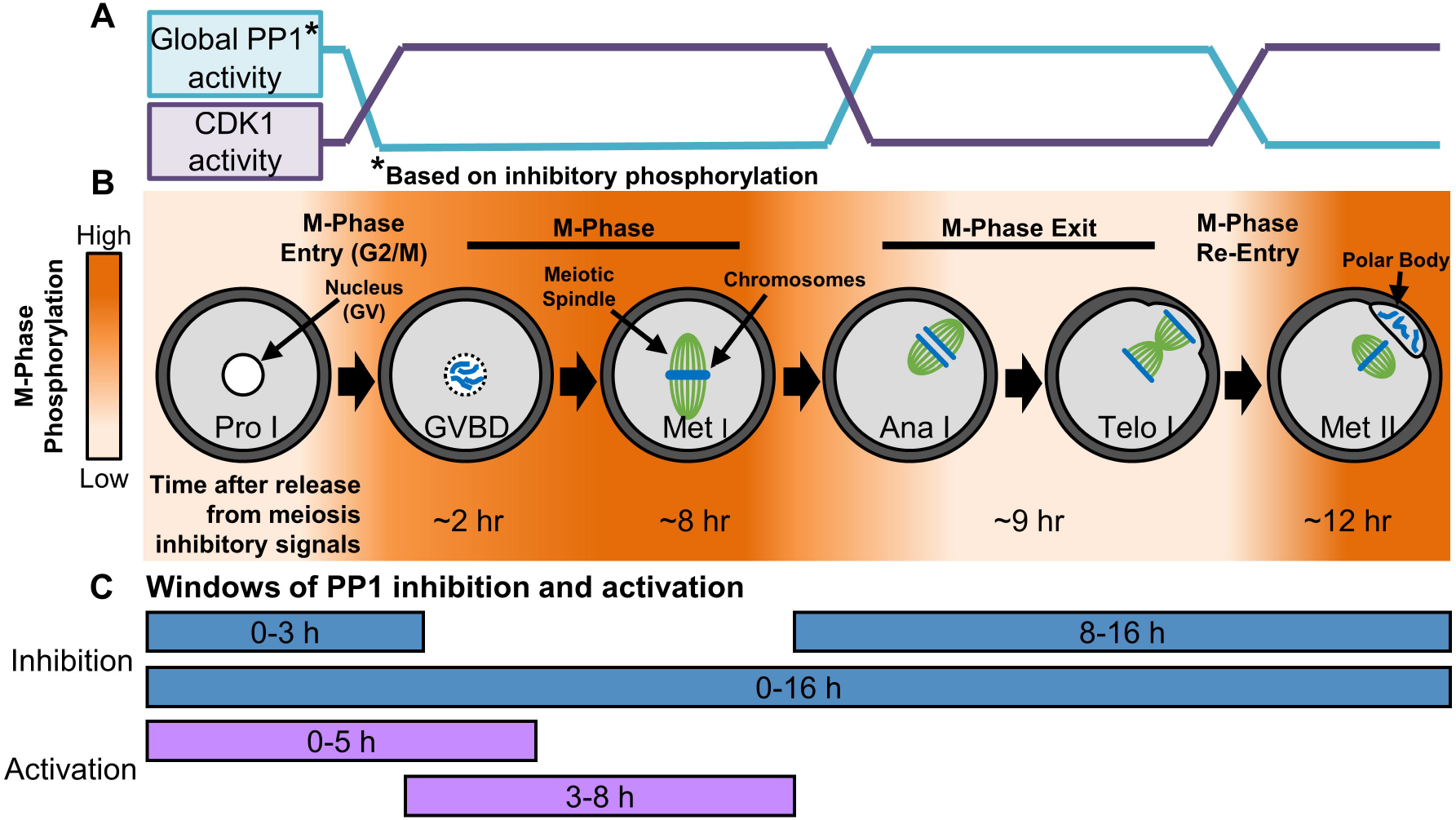
Phospho-dynamics and M-Phase regulation in oocyte meiosis. (A) Enzymatic activity of CDK1 and PP1 in mouse oocytes from prophase I to metaphase II, a process known as meiotic maturation. PP1 activity is shown as global activity based on the presence of PP1 inhibitory phosphorylation (pT320 on PP1cα). (B) Oocytes are arrested at prophase I (Pro I) until just prior to ovulation. Resumption of meiosis and entry into meiosis I M-Phase is equivalent to the G2/M transition in mitosis. Oocytes progress through nuclear envelope breakdown (NEBD) and metaphase I (Met I), then exit M-Phase. After telophase I (Telo I) completion, oocytes exit meiosis I, but bypass interphase and re-enter M-Phase, arresting at metaphase II (Met II). In mice, it takes ~12 hours to progress from prophase I to metaphase II (at which point oocytes arrest until fertilization), with the time needed to reach each stage noted in the figure. Mass changes in protein phosphorylation drive this M-Phase entry and exit (orange = high phosphorylation, white = low phosphorylation). (C) Experimental windows when PP1 activity is inhibited with tautomycetin (TMC) or activated with PDP-Nal.

PP1 and PP2A are critically important for mitotic and meiotic M-Phase progression and exit. In mitosis, PP1 and PP2A have unique temporal and spatial regulation, and work in concert to oppose CDK1 activity, preventing M-Phase entry and promoting metaphase-to-anaphase transition (Bancroft et al., 2020; Grallert et al., 2015; Ma et al., 2016; Rogers et al., 2016). In mammalian oocytes, inhibition of PP1/PP2A with reagents such as okadaic acid and Calyculin A causes major meiotic abnormalities, and oocyte-specific conditional loss of PP2A causes substantial defects in chromosome segregation and M-Phase progression (Alexandre et al., 1991; Gavin et al., 1991; Hu et al., 2014; Mailhes et al., 2003; Schwartz and Schultz, 1991; Smith et al., 1998; Swain et al., 2007; Swain et al., 2003; Tang et al., 2016; Wang et al., 2004). However, data also suggest that PP2A is not the only phosphatase that regulates meiosis in mammalian oocytes, and so we sought to elucidate PP1’s functions in this important cell type, building on evidence that PP1 has important roles in meiosis in *Drosophila*, *Caenorhabditis elegans*, starfish oocytes, and mammalian spermatogenesis (Hattersley et al., 2016; Oppedisano et al., 2002; Swartz et al., 2021; Wang et al., 2019a).

PP1 is a holoenzyme consisting of a catalytic subunit (PP1c) and one to two regulatory subunits (called PIPs, for PP1-interacting subunit, or RIPPOs, for regulatory interactors of protein phosphatase 1) (Bollen et al., 2010; Heroes et al., 2013). In mammals, there are three PP1c genes and hundreds of RIPPOs, resulting in hundreds of unique holoenzymes with distinctive functions and substrates (Bollen et al., 2010; Heroes et al., 2013). Various forms of PP1 holoenzymes allow PP1 activity to contribute to numerous processes in mitosis, including the G2/M transition, microtubule-kinetochore attachments, the spindle assembly checkpoint, and aspects of cytokinesis (Bancroft et al., 2020; Bhowmick et al., 2019; Capalbo et al., 2019; Cheng et al., 2000; Conti et al., 2019; Margolis et al., 2003; Smith et al., 2019; Winkler et al., 2015). Oscillations in phosphorylation of RIPPOs and PP1c, particularly phosphorylation of PP1c’s inhibitory site (pT320 on PP1cα and the equivalent sites in PP1cβ and PP1cγ), during M-Phase play a critical role in the formation of PP1 holoenzymes and PP1 activity, and ultimately mitotic M-Phase progression (Dohadwala et al., 1994; Kwon et al., 1997; Nasa et al., 2018). Notably, oscillations in RIPPO and PP1c phosphorylations in M-Phase of starfish and mammalian oocytes mirror those observed in mitotic cells (Adhikari et al., 2012; Dohadwala et al., 1994; Kwon et al., 1997; Swartz et al., 2021).

As noted above, exposure of mammalian oocytes to okadaic acid or Calyculin A, which inhibit PP1 and PP2A, leads to significant aberrations during *in vitro* meiotic maturation, including accelerated nuclear envelope breakdown (NEBD, G2/M transition), abnormal spindle morphology, meiosis I failure, and aneuploidy (Alexandre et al., 1991; Gavin et al., 1991; Mailhes et al., 2003; Schwartz and Schultz, 1991; Smith et al., 1998; Swain et al., 2007; Swain et al., 2003; Wang et al., 2004). However, only two studies have attempted to tease apart the functions of PP1 and PP2A, and these studies suggest that that PP1 is important for NEBD and chromosome condensation in mouse oocytes (Swain et al., 2007; Swain et al., 2003). To establish the roles that PP1 plays in mammalian oocyte M-Phase, we leveraged a small molecule approach to manipulate PP1 activity in opposition to its anticipated activity (e.g., inappropriately activating PP1 at G2/M) or preventing oscillations in PP1 activity (e.g., inhibiting PP1 for the duration of meiosis). It is worth noting that mammalian oocytes present a valuable and unique opportunity to unravel the functions of cell cycle regulatory proteins due to their relatively slow and synchronous progression through M-Phase of meiosis I. As a result of this synchronicity and extended M-Phase (7-10 h), mouse oocytes offer an exceptional model for the precise study of PP1 at discrete windows of time (e.g., G2/M transition only), without the need for potentially damaging synchronization methods. To leverage these strengths of the mouse oocyte system and to investigate PP1 roles with unprecedented control, mouse oocytes were exposed to the PP1 selective inhibitor tautomycetin (Mitsuhashi et al., 2001) and a recently developed PP1 activator PDP-Nal (Wang et al., 2019b) at discrete windows of meiosis (Fig. 1C). These studies reveal that alternation of normal PP1 activity fluctuations caused a range of severe meiotic defects, pointing to roles for PP1 in oocyte M-Phase and female fertility.

## Results

### Tautomycetin (TMC) selectively inhibits PP1 in mouse oocytes

Tautomycetin (TMC) is a PP1 selective inhibitor, with a ~40 fold higher affinity for PP1 over PP2A (Mitsuhashi et al., 2001). We first examined TMC’s selectivity for PP1 in mouse oocytes at 5 μM by assessing a PP1 substrate (phosphorylation of Histone H3 on Threonine 3; pH3T3) and a PP2A substrate (phosphorylation of AKT on Serine 473; pAKT(S473)). TMC was tested at 5 μM because this concentration of TMC causes a 90% reduction in PP1 activity with no impact on PP2A activity in COS-7 cells (Mitsuhashi et al., 2003). To assess how phosphorylation of these PP1 and PP2A substrates was affected by phosphatase inhibitors, oocytes were exposed to TMC or to Calyculin A, which is an inhibitor of both PP2A and PP1. Inhibitor treatments were done in time windows when phosphatase activity would be expected to be high, and thus phosphorylation of specific phosphatase substrates would be expected to be low. H3T3, a PP1 substrate, is dephosphorylated at M-Phase exit (Nguyen et al., 2014; Wang et al., 2016), with a significant decrease in pH3T3 observed at 9 h post-meiotic release (Fig. 2A). Treatment of oocytes with 5 μM TMC from 0 - 7 h or 0 - 9 h post-meiotic arrest release caused a significant increase in the PP1 substrate pH3T3 compared to vehicle control (Fig. 2B; 7 h, 23.1 ± 5.3% pH3T3 increase, p=0.0332; 9 h, 41.0 ± 4.9% pH3T3 increase, p=0.0002), confirming that TMC inhibits PP1. To validate the selectivity of TMC, immunofluorescence analysis of the PP2A substrate pAKT(S473) was performed. AKT(S473) has dynamic M-Phase phosphorylation (Han et al., 2006; Kalous et al., 2006), with pAKT(S473) significantly decreasing at 3 h post-meiotic arrest release (M-Phase entry / early M-Phase; Fig. 2C). Treatment of oocytes with 50 nM Calyculin A (CalA; PP1/PP2A inhibitor) from 2 - 3 h or 2 - 4 h post-meiotic arrest release caused a significant increase in phosphorylation of pAKT(S473) (Fig. 2D; 3 h, 91.5 ± 7.8% pAKT(S473) increase, p<0.0001; 4 h, 268.9 ± 7.7% pAKT(S473) increase, p<0.0001). In contrast, treatment of oocytes with 5 μM TMC in these same time windows did not cause a change in phosphorylation of the PP2A substrate pAKT(S473) (Fig. 2D; 3 h, p>0.9999; 4 h, p=0.4901). Taken together, these data highlight the selectivity of 5 μM TMC for PP1 in mouse oocytes.

**Figure 2:**
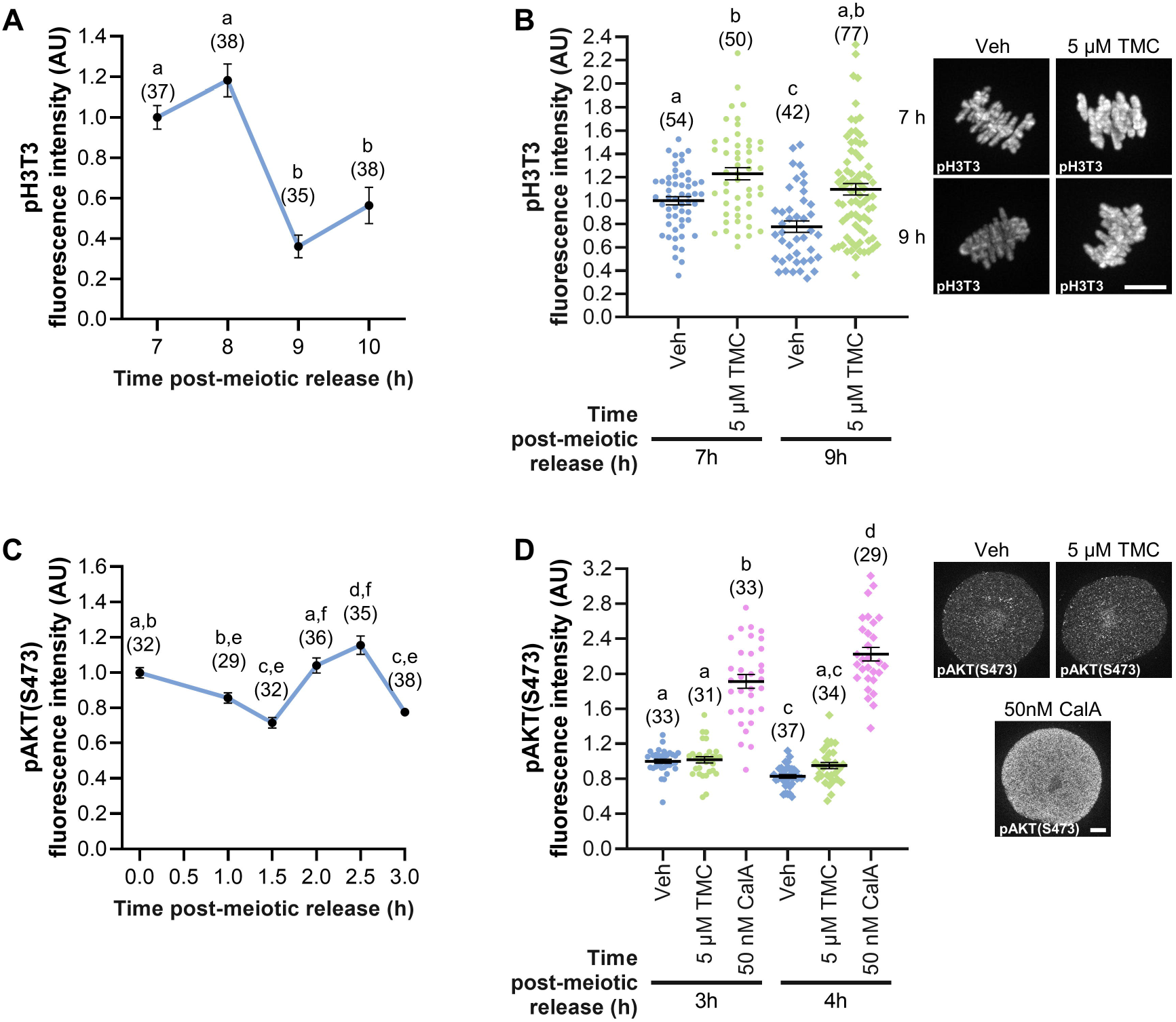
TMC increases PP1, but not PP2A, substrate phosphorlaytion. (A) Graphical comparison of pH3T3 fluorescence intensity over time in control, untreated oocytes. pH3T3 fluorescence signal was normalized to expression at 7 h post-meiotic arrest release. Different letters denote a significant difference between the groups (Kruskal-Wallis test with Dunn’s post hoc). n=35-38 oocytes over two replicates. (B) Graphical comparison of pH3T3 fluorescence intensity. Prophase I oocytes were released from meiotic arrest and cultured in medium containing TMC (5 μM) or vehicle control (0.1% DMSO) for 7 or 9 h. pH3T3 fluorescence signal was normalized to the signal in Veh oocytes at 7 h. Different letters denote a significant difference between the groups (Kruskal-Wallis test with Dunn’s post hoc). n=42-77 oocytes over two replicates. Representative images of oocyte pH3T3 fluorescence, scale = 10 μm. (C) Graphical comparison of pAKT(S473) fluorescence intensity over time in control, untreated oocytes. pAKT(S473) fluorescence signal was normalized to the signal at 0 h post-meiotic arrest release. Different letters denote a significant difference between the groups (ANOVA with Tukey’s post hoc). n=29-38 oocytes over two replicates. (B) Graphical comparison of pAKT(S473) fluorescence intensity. Prophase I oocytes were released from meiotic arrest and cultured for 2 h before exposure to medium containing TMC (5 μM), CalA (50 nM) or vehicle control (0.1% DMSO) for 1 h (3 h post-meiotic release) or 2 h (4 h post-meiotic release). pAKT(S473) fluorescence was normalized to expression in Veh oocytes at 3 h. Different letters denote a significant difference between the groups (Kruskal-Wallis test with Dunn’s post hoc). n=29-37 oocytes over two replicates. Representative images of oocyte pAKT(S473) fluorescence, scale = 10 μm. Line graphs show mean and SEM over time. Scatter dot plots show each individual oocyte fluorescence intensity plus mean and SEM.

### PP1 inhibition through meiotic maturation impairs M-Phase exit and reduces oocyte quality

Prior studies investigating the role of PP1 in mammalian oocytes used inhibitors that inhibit both PP1 and PP2A (e.g., okadaic acid and CalA), which complicated interpretation of data and prevented ascribing effects to one or the other phosphatase. For our initial assessment of the functions of PP1 during mouse oocyte meiosis, we exposed oocytes to TMC for the duration of *in vitro* maturation (IVM; 0-16 h; Fig. 3A), and assessed the ability of oocytes to progress through meiosis I and to metaphase II. TMC impaired M-Phase exit compared to vehicle control, as determined by the presence of a polar body (PB) by phase microscopy (Fig. 3B; Veh, 93.6% PB emission (PBE); 0.5 μM TMC 79.2% PBE, p=0.0216; 1 μM TMC 95.9% PBE, p=0.7154; 2.5 μM TMC 77.6% PBE, p=0.0119; 5 μM TMC 70.1% PBE, p<0.0001). Using immunofluorescence, we further characterized the meiotic stage of oocytes and assessed features of oocyte quality. Oocytes were scored as normal or abnormal, with classification as abnormal if a normal metaphase II DNA plate was not present after 16 h of IVM (examples of abnormalities observed in Fig. 3D-J’). PP1 inhibition resulted in a substantial increase in meiotic abnormalities in a dose-dependent manner compared to vehicle control (Fig. 3C-K; meiotic abnormalities: Veh 10.7% abnormal; 0.5 μM TMC 26.9% abnormal, p=0.1010; 1 μM TMC 16.1% abnormal, p=0.5114; 2.5 μM TMC 34.3% abnormal, p=0.0131; and 5 μM TMC 88.0% abnormal, p<0.0001). These abnormalities ranged from an oocyte being arrested in metaphase I (Fig. 3D) to two spindles with no PB (Fig. 3G), with increasing doses of TMC leading to more severe defects.

**Figure 3:**
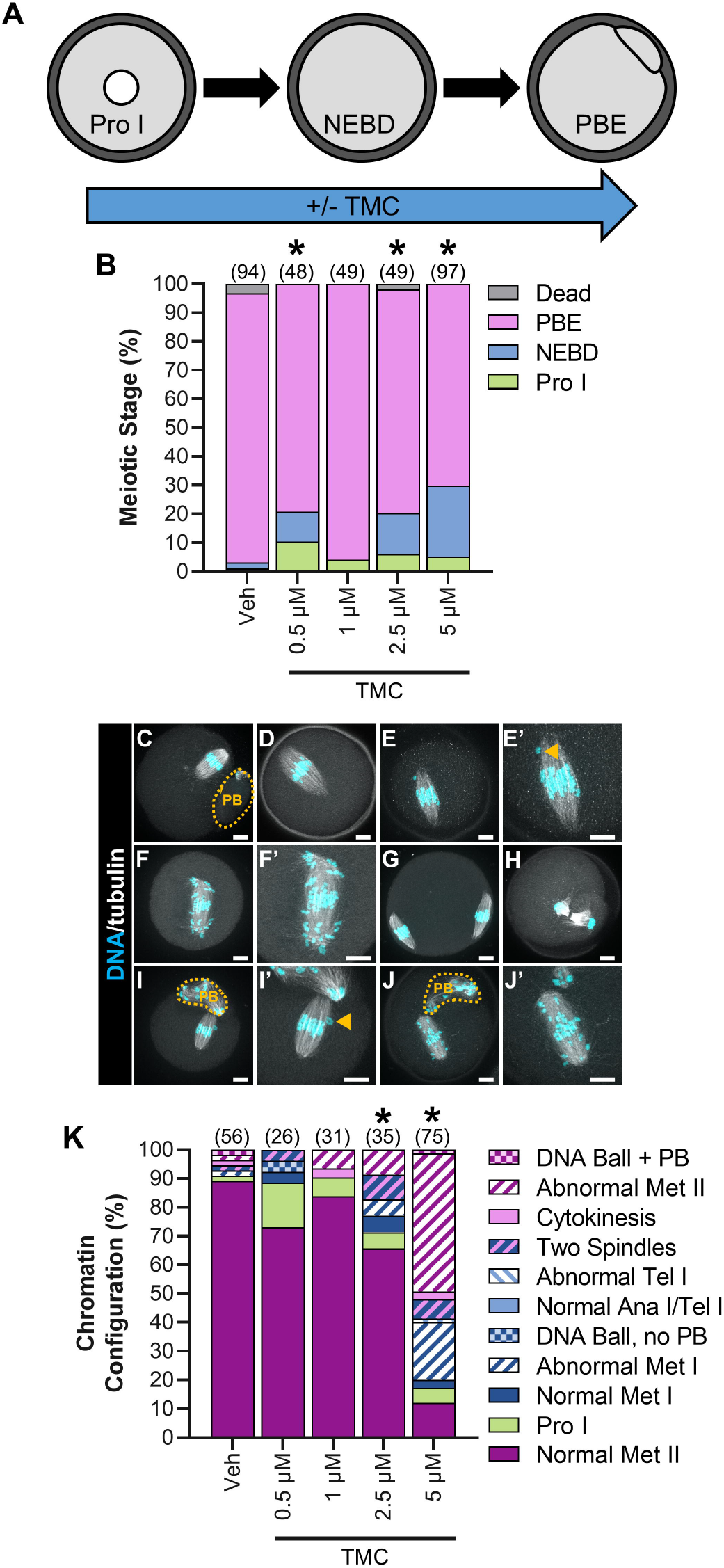
PP1 inhibition for the duration of oocyte meiotic maturation reduces meiosis I exit in oocytes and causes meiotic abnormalities. (A) Schematic representation of the experimental design. Prophase I oocytes were cultured in medium lacking meiotic arrest reagents and containing TMC (0.5-5 μM) or vehicle control (0.1% DMSO) for 16 h. (B) Graphical representation of meiotic stage based on phase microscopy at 16 h into culture. Bar graph shows the percentage of oocytes at each meiotic stage. n=48-97 oocytes over 3-5 replicates. * denotes a significant difference from Veh control oocytes (Fisher’s exact test). (C-J) Representative images of the oocyte phenotypes observed, apostrophe denotes zoomed-in image, scale bar = 10 μM. Orange dashed line denotes the PB boundary, and orange arrow points to single misaligned chromosomes. (C) Normal metaphase II oocyte. Oocytes that underwent NEBD but had no PB with a normal (D) metaphase I spindle, with (E and E’) mild chromosome misalignment, with (F and F’) severe chromosome misalignment, or with (G) two spindles. (H) Oocyte at cytokinesis. Metaphase II oocytes, with (I and I’) mild chromosome misalignment, or with (J and J’) severe chromosome misalignment. Oocytes with DNA ball, ± PB refers to very condensed chromatin in the oocyte, with no obvious meiotic spindle (not shown) (K) Graphical representation of the meiotic stage and DNA morphology of oocytes at 16 h into culture. Solid colors represent normal DNA morphology for the meiotic stage, and hatched colors represent abnormal DNA morphology for the meiotic stage. n=26-75 oocytes over 3-5 replicates. * denotes a significant difference from Veh control oocytes (Fisher’s exact test).

We attempted similar experiments for the duration of IVM with a reagent to manipulate PP1 to an activated state, treating oocytes with the recently developed PP1 activating peptide PDP-Nal (Wang et al., 2019b). PDP-Nal stimulates PP1 activity by uncoupling the PP1 catalytic subunit from RIPPO regulation. However, exposure of oocytes to PDP-Nal for the duration of IVM (0-16 h) resulted in 100% of oocytes dying. Taken together, these data highlight that oscillations in PP1 activity is essential for normal meiotic progression.

### Reduced PP1 activity drives G2/M transition in oocytes

In mitosis, oscillations in PP1 activity are essential for M-Phase progression, with decreased PP1 activity driving G2/M transition and increased PP1 activity being crucial for metaphase/anaphase transition (Moura and Conde, 2019). This cyclic PP1 activity is driven in part by a CDK1-mediated inhibitory phosphorylation of PP1c (pPP1cα(T320) or equivalent sites in PP1cβ and PP1cγ) (Dohadwala et al., 1994; Kwon et al., 1997). The kinetics of PP1 inhibitory phosphorylation follows an identical pattern in starfish and mouse oocytes, suggesting similar fluctuations in PP1 activity during oocyte meiosis (Adhikari et al., 2012; Swartz et al., 2021). Therefore, we sought to determine if a reduction in PP1 activity is critical for meiotic resumption by culturing oocytes in PDP-Nal during the G2/M transition (0-5 h post-meiotic arrest release; Fig. 4A). Oocytes were released from prophase I arrest into increasing concentrations of PDP-Nal, the negative control peptide (PDPm-Nal; which has no PP1c binding), or standard culture medium (Ctrl). G2/M transition, as determined by NEBD, was hampered by PP1 activation in a dose-dependent manner compared to PDPm-Nal treatment (Fig. 4B; NEBD at 5 h: PDPm-Nal 86.7%; Ctrl 86.8%, p >0.9999; 2.5 μM PDP-Nal 74.0%, p=0.0031; 5 μM PDP-Nal 50.3%, p<0.0001; 7.5 μM PDP-Nal 35.7%, p<0.0001; 10 μM PDP-Nal 19.3%, p<0.0001). Moreover, PP1 activation significantly reduced oocyte survival (Fig. 4C; Dead oocytes at 5 h: PDPm-Nal 2.3%; Ctrl 0.6%, p=0.2147; 2.5 μM PDP-Nal 12.4%, p=0.0003; 5 μM PDP-Nal 39.5%, p<0.0001; 7.5 μM PDP-Nal 50.0%, p<0.0001; 10 μM PDP-Nal 66.9%, p<0.0001). Critically, the majority of oocyte death was observed in prophase I oocytes, rather than oocytes that had resumed meiosis (Fig. 4D; % of oocytes that died at prophase I: 2.5 μM 85.7%; 5 μM 83.5%; 7.5 μM 85.7%; 10 μM 91.7%).

**Figure 4:**
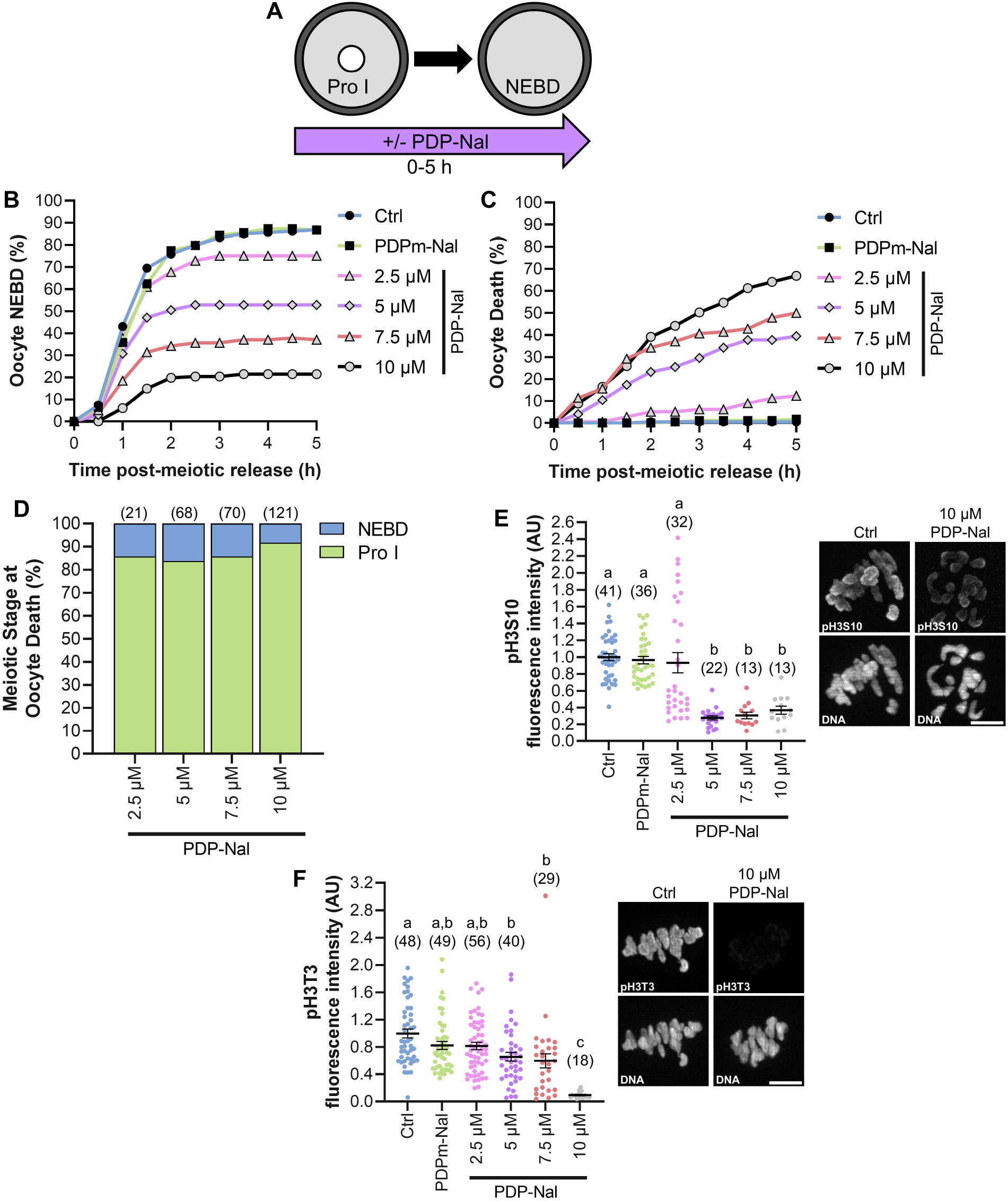
PP1 activation impairs meiotic resumption. (A) Schematic representation of the experimental design. Prophase I oocytes were released from meiotic arrest into culture medium containing PDP-Nal (2.5-10 μM), 10 μM PDPm-Nal, or standard culture medium (Ctrl; 0 μM PDP-Nal). NEBD was monitored to track meiotic resumption. (B-C) Graphical representation of the percentage of oocytes to (B) resume meiosis (NEBD) or (C) die over time. n=140-181 oocytes over 7-9 replicates. (D) Graphical representation of meiotic stage at oocytes death; n=21-121 oocytes over 7-9 replicates. (E) Graphical comparison of pH3S10 fluorescence intensity. pH3S10 fluorescence was normalized to expression in Ctrl oocytes. Different letters denote a significant difference between the groups (Kruskal-Wallis test with Dunn’s post hoc). n=13-41 oocytes over 2 replicates. Representative images of oocyte pH3S10 fluorescence, scale = 10 μm. (F) Graphical comparison of pH3T3 fluorescence intensity. pH3T3 fluorescence was normalized to expression in Ctrl oocytes. Different letters denote a significant difference between the groups (Kruskal-Wallis test with Dunn’s post hoc). n=18-56 oocytes over 3-4 replicates. Representative images of oocyte pH3T3 fluorescence, scale = 10 μm. Scatter dot plots show each individual oocyte fluorescence intensity plus mean and SEM.

To confirm that PP1 was activated, we investigated the phosphorylation status of two well characterized PP1 substrates, Histone H3 on Threonine 3 (pH3T3) and on Serine 10 (pH3S10). pH3S10 was significantly decreased with 5, 7.5 and 10 μM PDP-Nal treatment compared to PDPm-Nal (Fig. 4E; 2.5 μM, 3.2 ± 12% pH3S10 loss, p > 0.9999; 5 μM, 71.1 ± 2.3% pH3S10 loss, p<0.0001; 7.5 μM, 68.2 ± 3.8% pH3S10 loss, p<0.0001; 10 μM, 61.2 ± 1.7% pH3S10 loss, p=0.0001). PP1 activation also caused a reduction of H3T3 phosphorylation, but this was only significant with the highest PDP-Nal concentration (10 μM) compared to PDPm-Nal oocytes (Fig. 4F; 2.5 μM, 0.9 ± 5.2% pH3T3 loss, p>0.9999; 5 μM, 20.1 ± 6.6% pH3T3 loss, p>0.9999; 7.5 μM, 27.5 ± 1.1% pH3T3 loss, p=0.2299; 10 μM, 88.6 ± 1.0% pH3T3 loss, p<0.0001).

To further define the role of PP1 activity at G2/M transition, we evaluated inappropriate exit from prophase I arrest, in conjunction with TMC treatment. This assay employed a modification of culture conditions used to maintain robust prophase I arrest for extended periods (≥ 16 h), culturing oocytes in 2.5 μM of the phosphodiesterase 3 inhibitor milrinone (Stein and Schindler, 2011; Tsafriri et al., 1996). To evaluate the effects of inhibiting PP1 on the maintenance of prophase I arrest, we used culture conditions that allow a portion of control oocytes to undergo the G2/M transition (NEBD) by culturing oocytes in a low concentration of milrinone (0.2 μM; Fig. 5A). With this lower concentration of milrinone (0.2 μM), 36.8% of Veh-treated oocytes were at prophase I arrest after 16 h of culture, compared to 6.2% of IVM control oocytes (no milrinone; p<0.0001). Significantly more TMC-treated oocytes (2.5 μM TMC) underwent the G2/M transition (NEBD) and resumed meiosis as compared to Veh-treated, as reflected by a smaller percentage of oocytes in these treatment groups being at prophase I after 16 h (Fig. 5B; Veh 36.8%; 0.5 μM TMC 26.4%, p=0.1826; 1 μM TMC 24.5%, p=0.1288; 2.5 μM TMC 16.2%, p=0.0018; 5 μM TMC 25.7%, p=0.0566). However, despite an increase in G2/M transition with TMC treatment, the percentage of oocytes to exit M-Phase was significantly reduced compared to vehicle control oocytes (Fig. 5B; % of oocytes arrested at NEBD at 16 h: Veh 1.3%; 0.5 μM TMC 7.5%, p=0.0399; 1 μM TMC 3.8%, p=0.2753; 2.5 μM TMC 29.7%, p<0.0001; 5 μM TMC 31.6%, p<0.0001).

**Figure 5:**
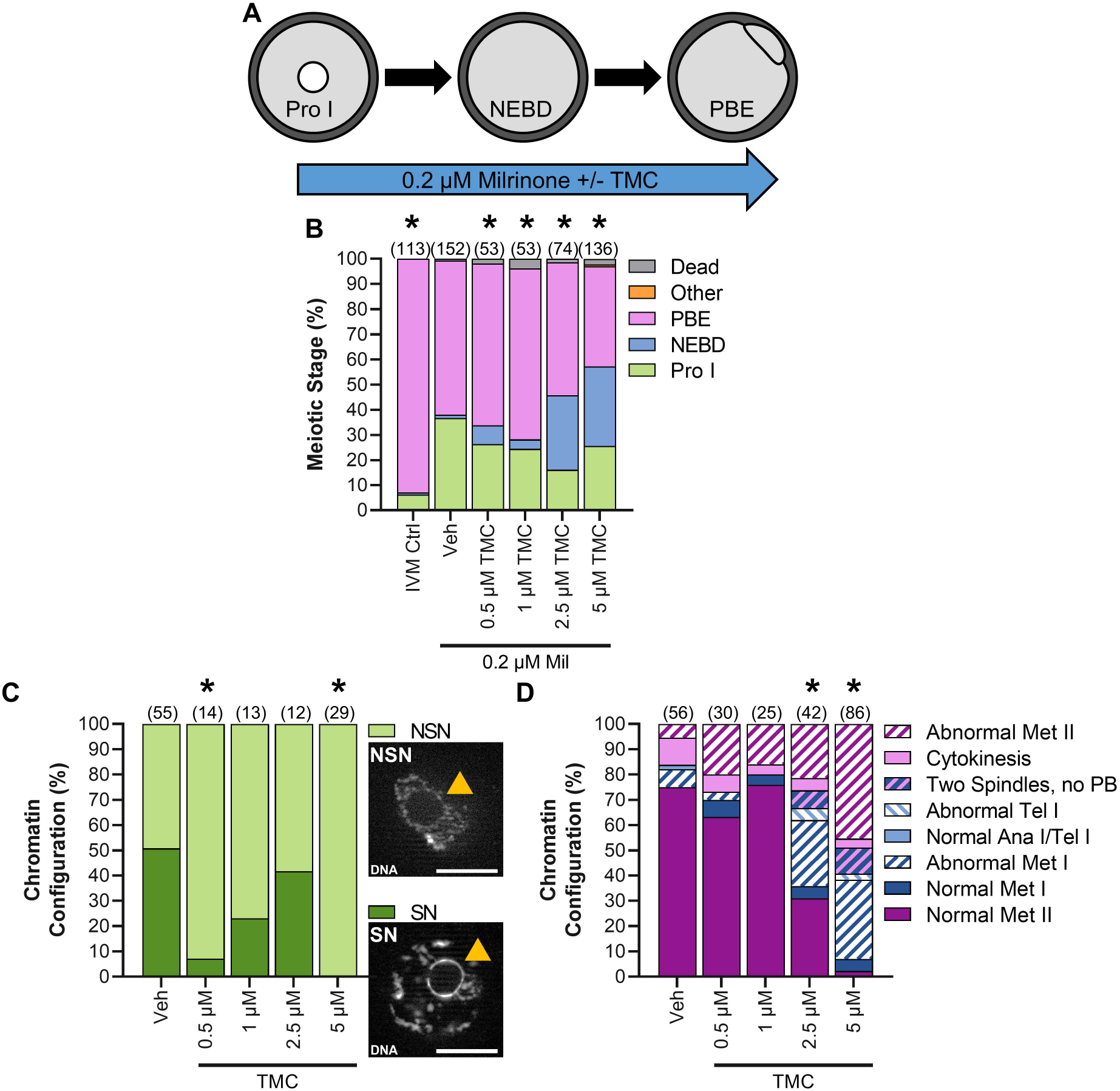
PP1 inhibition enhances NEBD, reduces M-Phase exit, and causes meiotic abnormalities in oocytes in culture conditions that allow partial maintenance of prophase I arrest. (A) Schematic representation of the experimental design. Prophase I oocytes were washed into medium containing 0.2 μM milrinone and increasing amounts of TMC (0.5-5 μM) or vehicle control (0.1% DMSO) for 16 h; under these culture conditions, ~37% of oocytes will undergo the G2/M transition, exiting from prophase I arrest. (B) Graphical representation of meiotic stage based on phase microscopy at 16 h into culture. IVM Ctrl oocytes were cultured in milrinone-free medium. Bar graph shows the percentage of oocytes at each meiotic stage. n=53-152 oocytes over 3-6 replicates. * denotes a significant difference from Veh control oocytes (Fisher’s exact test). (C) Graphical representation and images of DNA morphology in prophase I oocytes at 16 h into culture. Orange arrow points to the nucleolus and scale bar = 10 μM. n=12-55 oocytes over 3-6 replicates. * denotes a significant difference from Veh control oocytes (Fisher’s exact test). (D) Graphical representation of the meiotic stage and DNA morphology of oocytes at 16 h into culture. Solid colors represent normal DNA morphology for the meiotic stage, and hatched colors represent abnormal DNA morphology for the meiotic stage. n=25-86 oocytes over 3-6 replicates. * denotes a significant difference from Veh control oocytes (Fisher’s exact test).

Additionally, two other effects of TMC treatment in these experiments were observed. First, the chromatin configuration of prophase I oocytes was altered by TMC treatment. Chromatin configuration in prophase I oocytes is characterized by the extent of condensation of the DNA around the nucleolus, with two categories: surrounded nucleolus (SN) with a ring of condensed DNA around the nucleolus, and non-surrounded nucleolus (NSN) with uncondensed DNA (Fig. 5C). The transition from NSN to SN is correlated with acquisition of meiotic competence (ability of an oocyte to reach metaphase II) and with developmental potential (ability of a fertilized oocyte to support pre-implantation embryo development), and NSN oocytes display lower meiotic competence and developmental potential (Bellone et al., 2009; Zuccotti et al., 2002). Veh-treated oocytes had equal numbers of SN and NSN prophase I oocytes. In contrast, the majority of prophase I oocytes treated with 0.5 μM and 5 μM TMC had NSN configuration (Fig. 5C; Veh, 49.1% NSN; 0.5 μM TMC 92.9% NSN, p=0.0049; 5 μM TMC 100% NSN, p<0.0001). A similar trend towards increased NSN configuration was also observed with 1 μM and 2.5 μM TMC, although this did not reach significance, potentially due to low numbers of prophase I oocytes in these experimental groups (Fig. 5C; 1 μM TMC 76.9% NSN, p=0.1199; 2.5 μM TMC 58.3% NSN, p=0.7516). The second effect of TMC treatment was observed in oocytes that had resumed meiosis. TMC treatment resulted in a dose-dependent increase in meiotic abnormalities, similar to those observed with 16 h TMC treatment under standard IVM conditions (Fig. 5D; meiotic abnormalities: Veh 25% abnormal; 0.5 μM TMC 36.7% abnormal, p=0.3208; 1 μM TMC 24% abnormal, p>0.9999; 2.5 μM TMC 69% abnormal, p<0.0001; and 5 μM TMC 97.7% abnormal, p<0.0001). Taken together these results confirm that reducing PP1 activity is central for G2/M transition in oocytes.

### PP1 activation during M-Phase reduces oocyte survival and alters chromosome shape

Having established that inhibition of PP1 is essential for the G2/M transition, we next sought to determine the effects of premature PP1 activation during M-Phase. Oocytes were exposed to increasing concentrations of PDP-Nal from prometaphase (3 h post-meiotic arrest release) to metaphase I (8 h post-meiotic arrest release; Fig. 6A), when PP1 activity is low. To confirm that PP1 was activated, we investigated pH3T3 and pH3S10 abundance. At metaphase I (8 h post-meiotic arrest release), pH3T3 fluorescence in oocytes treated with 5 μM, 7.5 μM and 10 μM PDP-Nal was significantly decreased compared to PDPm-Nal-treated oocytes (Fig. 6B; 2.5 μM, 24.1 ± 4.7% pH3T3 loss, p= 0.3133; 5 μM, 43.1 ± 4.7% pH3T3 loss, p<0.0001; 7.5 μM, 44.8 ± 5.1% pH3T3 loss, p<0.0001; 10 μM, 46.0 ± 4.9% pH3T3 loss, p<0.0001). A more modest loss of pH3S10 was observed, with 5 and 7.5 μM PDP-Nal treatment significantly different compared to PDPm-Nal (Fig. 6C; 2.5 μM, 9.2 ± 4.2% pH3S10 loss, p>0.9999; 5 μM, 26.1 ± 7.8% pH3S10 loss, p=0.0057; 7.5 μM, 45.6 ± 5.0% pH3S10 loss, p<0.0001; 10 μM, 24.0 ± 6.0% pH3S10 loss, p=0.0647). To evaluate if we could achieve a higher extent of PP1 activation (and potentially lower pH3T3 and pH3S10 fluorescence signals at these time points), we tested two higher doses of PDP-Nal, 20 and 40 μM, which are required for sufficient PP1 activation in mitotic cells (Rogers et al., 2016; Wang et al., 2019b). However, treating oocytes with these higher doses of PDP-Nal led to 100% cell death (Fig. 6D; Dead oocytes at 8 h: 20 μM PDP-Nal 100% dead, p<0.0001; 40 μM PDP-Nal 100% dead, p<0.0001). Thus, subsequent experiments used PDP-Nal at 2.5, 5.0, 7.5, and 10 μM.

**Figure 6:**
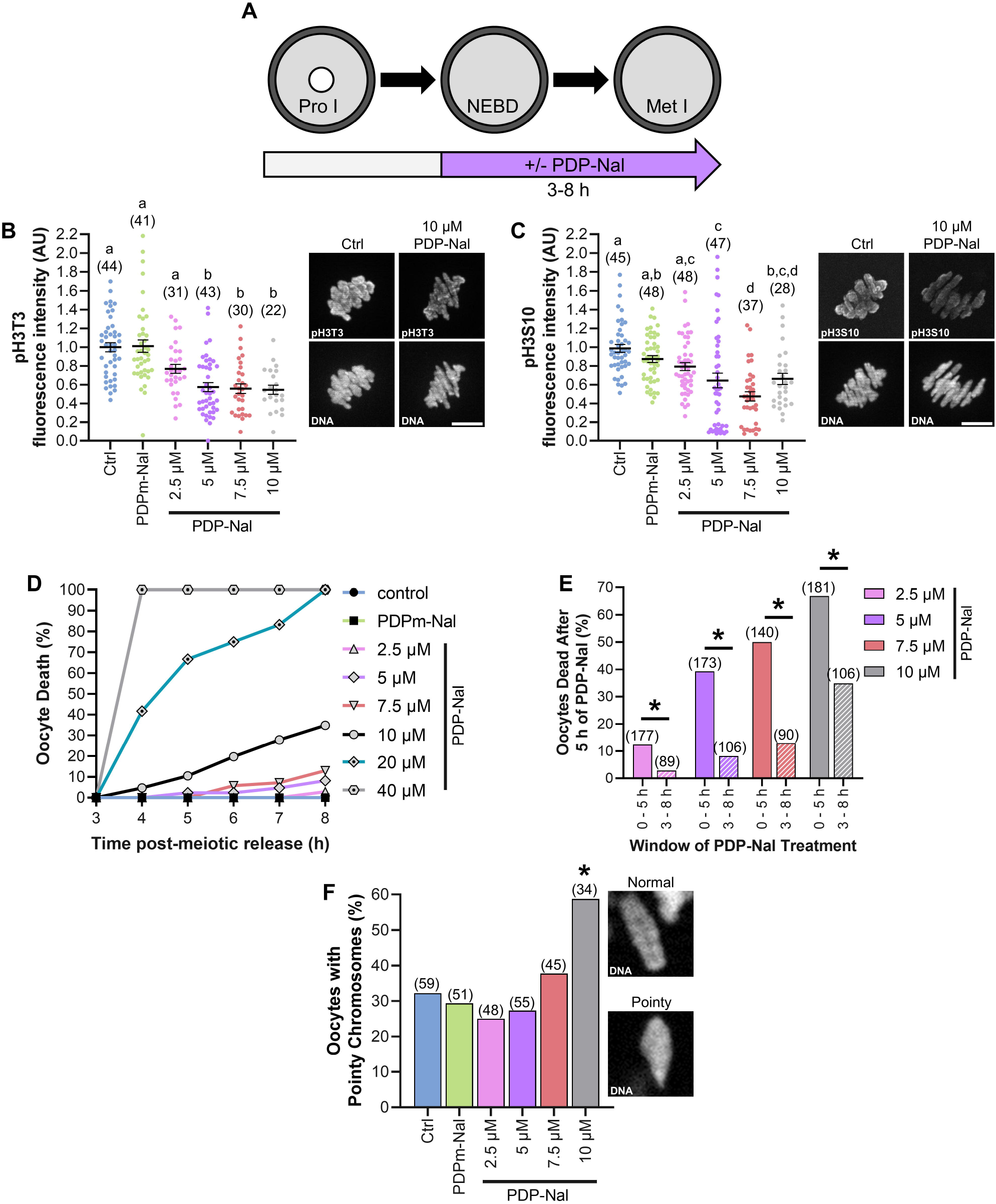
PP1 activation from prometaphase to metaphase of meiosis I decreases oocyte survival and alters chromosome morphology. (A) Schematic representation of the experimental design. Prophase I oocytes were released from meiotic arrest and cultured for 3 h before treatment with increasing concentrations of PDP-Nal (2.5-40 μM), PDPm-Nal (10 μM) or standard culture medium (Ctrl; 0 μM PDP-Nal). (B) Graphical comparison of pH3T3 fluorescence intensity. pH3T3 fluorescence was normalized to expression in Ctrl oocytes. Different letters denote a significant difference between the groups (Kruskal-Wallis test with Dunn’s post hoc). n=22-41 oocytes over three replicates. Representative images of oocyte pH3T3 fluorescence, scale = 10 μm. (C) Graphical comparison of pH3S10 fluorescence intensity. pH3S10 fluorescence was normalized to expression in Ctrl oocytes. Different letters denote a significant difference between the groups (Kruskal-Wallis test with Dunn’s post hoc). n=28-48 oocytes over three replicates. Representative images of oocyte pH3S10 fluorescence, scale = 10 μm. (D) Graphical representation of the percentage of oocyte death over time. n=29-106 oocytes over 2-5 replicates. (E) Graphical representation of percentage of oocytes dead with 5 h of PDP-Nal treatment at G/2M (0-5 h; solid color) or during meiosis I (3-8 h; hashed color). * denotes a significant difference between 0-5 h and 3-8 h PDP-Nal treated oocytes (Fisher’s exact test) (F) Graphical representation of the percentage of oocytes with at least one pointy chromosome, with sample images of DAPI staining showing chromosome morphology. n=34-59 oocytes over five replicates. * denotes a significant difference from PDPm-Nal control oocytes (Fisher’s exact test). Scatter dot plots show each individual oocyte fluorescence intensity plus mean and SEM.

PP1 activation for this five-hour period from prometaphase to metaphase I caused a dose-dependent increase in oocyte death, compared to no death observed in PDPm-Nal-treated oocytes (Fig. 6D; % of oocytes dead at 8 h: PDPm-Nal 0%; Ctrl 0%, p>0.9999; 2.5 μM PDP-Nal 3.4%, p=0.0963; 5 μM PDP-Nal 7.5%, p=0.0068; 7.5 μM PDP-Nal 13.3%, p<0.0001; 10 μM PDP-Nal 33.0%, p<0.0001). We also compared the extent of oocyte death observed with 5 h of PDP-Nal treatment from prometaphase to metaphase I and during the G2/M transition (0-5 h post-meiotic arrest release). Surprisingly, PP1 activation for this five-hour period during M-Phase (3-8 h post-meiotic arrest release) was less deleterious than PP1 activation for five hours during the G2/M transition (0-5 h post-meiotic arrest release). Significantly higher numbers of oocytes survived PDP-Nal treatment during M-Phase than during the G2/M transition (Fig. 6E; 2.5 μM: 12.4% (0-5 h) vs. 2.9% (3-8 h) dead, p=0.0151; 5 μM: 39.3% (0-5 h) vs. 8.2% (3-8 h) dead, p<0.0001; 7.5 μM: 50% (0-5 h) vs. 13% (3-8 h) dead, p<0.0001;10 μM: 66.9% (0-5 h) vs. 34.9% (3-8 h) dead, p<0.0001).

PP1 has roles in mitotic chromosome alignment and actin regulation (Canals et al., 2012; Carmody et al., 2008; Vallardi et al., 2017; Vallée et al., 2018). In oocytes, actin is essential for migration of the meiotic spindle from the center of the oocyte to the cortex, and thus for asymmetrical cytokinesis (Evans and Robinson, 2011; Uraji et al., 2018). Therefore, chromosome alignment on the metaphase I plate and the distance of the metaphase I plate to the cortex was assessed in oocytes following PP1 activation from prometaphase to metaphase I (3 - 8 h post-meiotic release). PP1 activation did not appear to affect these features of the metaphase I spindle (Supp Fig. S1). However, PP1 activation did alter the shape of individual chromosomes, with a shift from normal morphology of chromosomes with rounded ends, to chromosomes with pointy ends observed at 10 μM PDP-Nal treatment compared to PDPm-Nal (Fig. 6F; Oocytes with at least one pointy chromosome: PDPm-Nal 29.4%; Ctrl 32.2%, p=0.8372; 2.5 μM PDP-Nal 25.0%, p=0.6577; 5 μM PDP-Nal 27.3%, p=0.8322; 7.5 μM PDP-Nal 37.8%, p=0.3964; 10 μM PDP-Nal 58.8%, p=0.0127). Taken together, these results show that reduced PP1 activity during M-Phase plays a key role in oocyte survival and chromosome architecture.

### PP1 inhibition during M-Phase slows the timing of meiosis I exit

We next sought to determine the effect of PP1 inhibition during M-Phase, given that PP1 activity would be anticipated to be important for meiosis I exit. As our studies above found that TMC treatment for the duration of meiotic maturation impaired meiosis I exit (Fig. 3), we treated oocytes with 5 μM TMC from 0 - 7 h or 0 - 9 h post-meiotic arrest release (Fig. 7A). These timepoints were selected because analyses of chromatin appearance (Fig. 7B) and phosphorylation of the PP1 substrate H3T3 in control untreated oocytes (Fig. 2A and Supp Fig. S2) revealed that the 7-hour timepoint corresponded to late prometaphase / early metaphase I with relatively low global PP1 activity, and the 9-hour timepoint corresponded to the start of M-Phase exit and relatively high global PP1 activity.

**Figure 7:**
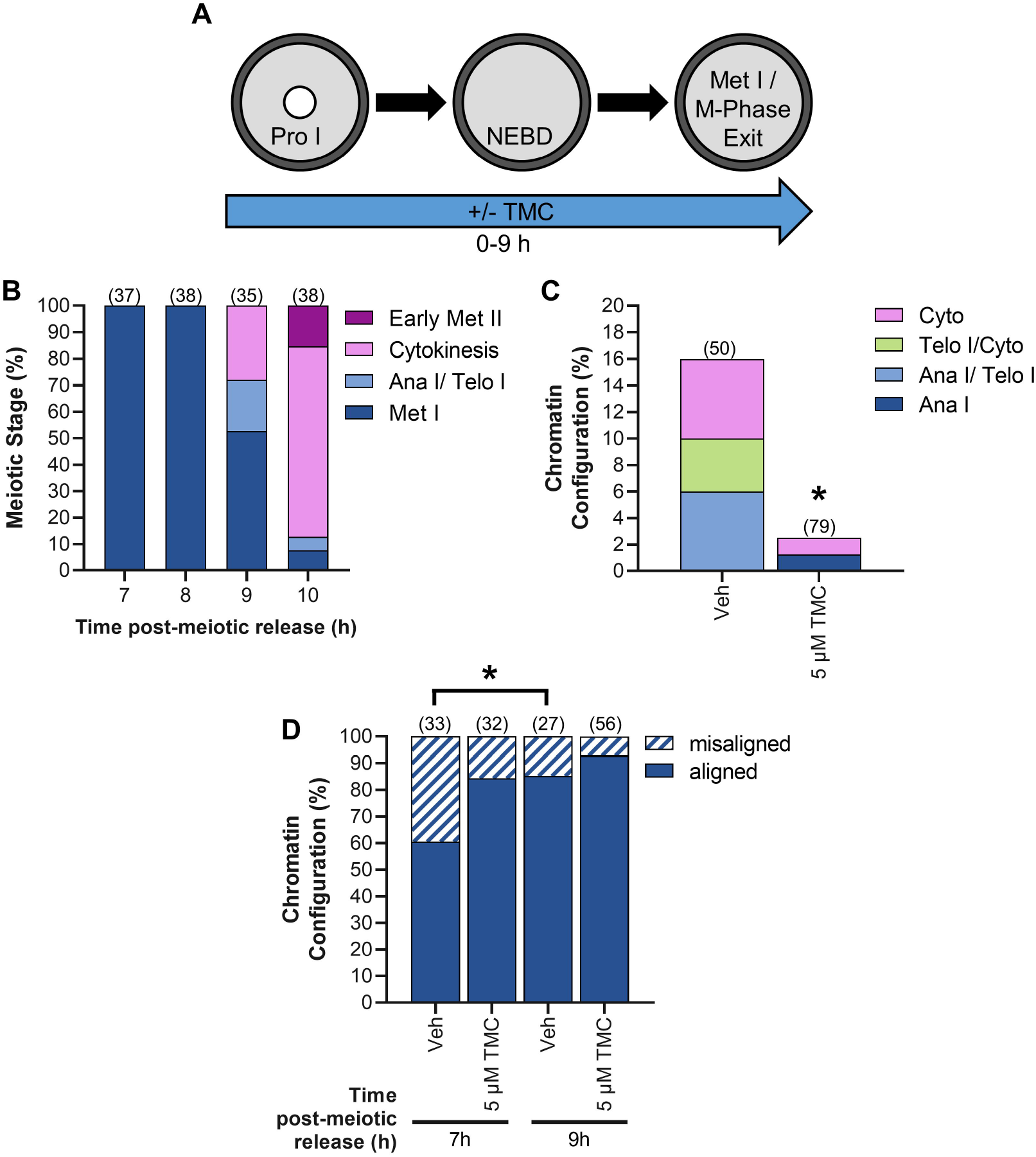
PP1 inhibition during meiosis I impairs M-Phase exit. (A) Schematic representation of the experimental design. Prophase I oocytes were released from meiotic arrest and cultured in medium containing TMC (5 μM) or vehicle control (0.1% DMSO) for 7 or 9 h. (B) Graphical representation of meiotic stages in control untreated oocytes at 7, 8, 9, and 10 h post-meiotic arrest release. n=35-38 oocytes over two replicates. (C) Graphical representation of stages of meiotic exit at 9 h post-meiotic arrest release. * denotes a significant difference from Veh control oocytes (Fisher’s exact test). n=50-79 oocytes over two replicates. (D) Graphical representation of chromosome alignment in metaphase I oocytes at 7 or 9 h post-meiotic arrest release. * denotes a significant difference (Fisher’s exact test). n=27-56 oocytes over two replicates.

At 7 h, 100% of vehicle control and TMC-treated oocytes were at metaphase I (Veh, n=54 oocytes; TMC, n=50 oocytes). By 9 h, 16% of vehicle control oocytes were beginning to exit meiosis I, classified as anaphase I to cytokinesis, whereas only 2.5% TMC-treated oocytes reached this stage of M-Phase exit (Fig. 7C; p=0.0134). We also examined the metaphase I plate of oocytes for chromosome alignment and distance from the cortex (as illustrated in Supp Fig. 2B-C). At 7 h, 60.6% of Veh-treated oocytes and 85.2% of TMC-treated oocytes had an aligned metaphase I plate (just outside of significance at p=0.0514). At 9 h, 84.4% of Veh-treated had an aligned metaphase I plate (with this increase from 7 h being statistically significant, p=0.0463), and 92.9% of TMC-treated oocytes had an aligned metaphase I plate (Fig. 7D; p=0.2764 as compared to 7 h). TMC treatment did not appear to affect metaphase spindle migration to the cortex, with no difference in the distance of the metaphase plate to the cortex between the vehicle control and TMC-treated groups (Supp Fig. S3).

### Reduced PP1 activity from metaphase I onwards impairs M-Phase exit, but has no detectable effects on oocyte quality

Our studies noted above (Fig. 3K) showed that TMC treatment for the duration of meiotic maturation significantly impaired meiosis I exit, with 22.7% of 5 μM TMC-treated oocytes arrested at metaphase I at 16 h IVM compared to 1.8% of Veh-treated oocytes. Similarly, PP1 inhibition in mitotic cells caused arrest at metaphase, with PP1 found to have essential roles at the metaphase/anaphase transition (Winkler et al., 2015). To determine if the metaphase I arrest phenotype we observed resulted from a loss of PP1 activity specifically at metaphase I/anaphase I transition, oocytes were treated with TMC in this precise time window from metaphase I onward (8 - 16 h post-meiotic arrest release; Fig. 8A). TMC treatment during this time of meiosis reduced M-Phase exit compared to vehicle control, as determined by the presence of a polar body by phase microscopy (Fig. 8B; Veh, 100% PBE; 0.5 μM TMC 97.3% PBE, p.0.9999; 1 μM TMC 96.2% PBE, p=0.4955; 2.5 μM TMC 87.6% PBE, p=0.0267; 5 μM TMC 84.7% PBE, p<0.0001). However, the extent of impaired M-Phase exit in these experiments was substantially less than that observed with TMC treatment for the duration of meiosis (0 - 16 h post-meiotic arrest release). Furthermore, TMC treatment during this time of meiosis did not cause the meiotic abnormalities observed with the longer duration treatment (Fig. 8C; meiotic abnormalities: Veh = 15.2% abnormal; 0.5 μM TMC = 8.8% abnormal, p=0.4763; 1 μM TMC = 12.1% abnormal, p>0.9999; 2.5 μM TMC = 22.5% abnormal, p=0.5542; and 5 μM TMC 26.5% abnormal, p=0.3689). These results point to PP1 having critical roles earlier in meiosis.

**Figure 8:**
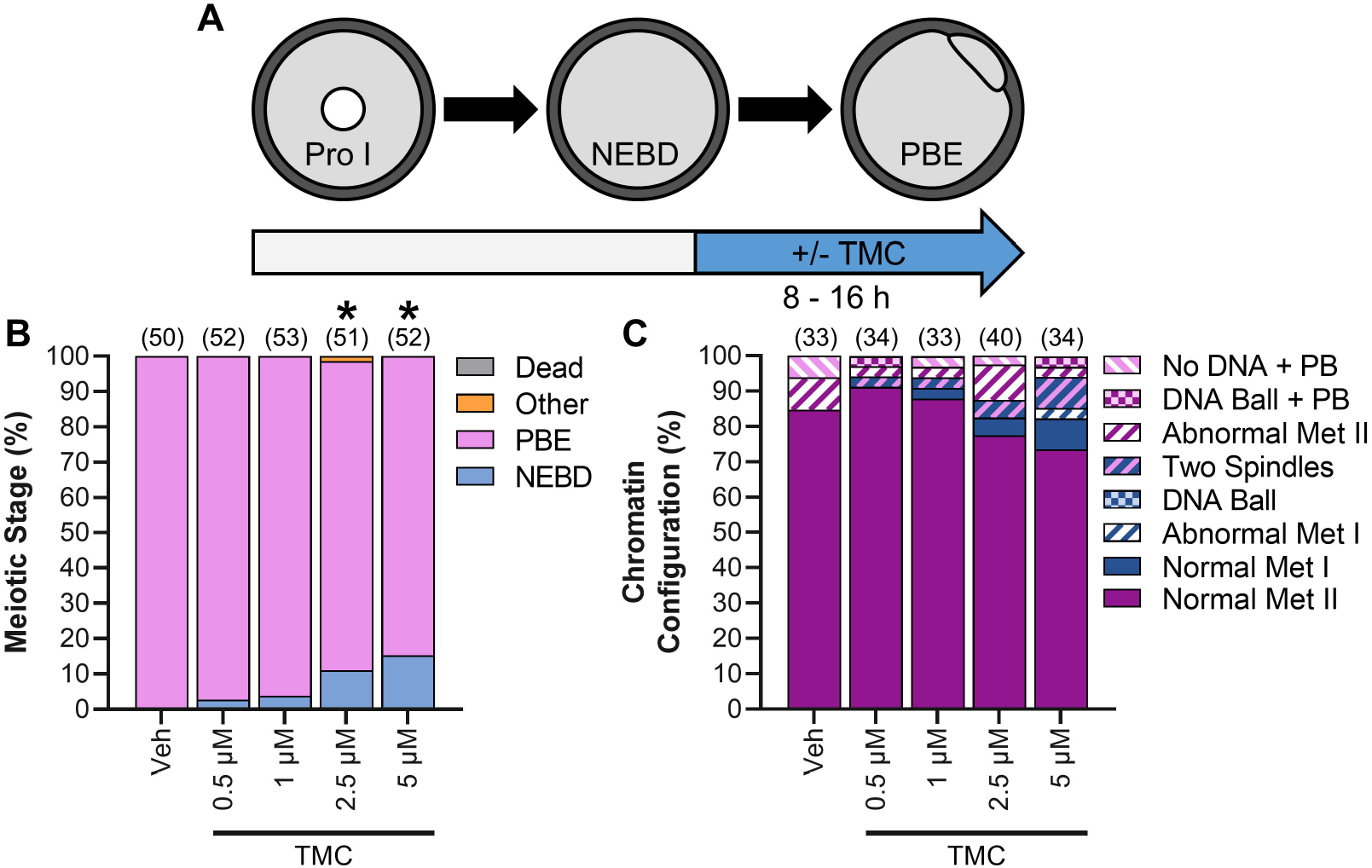
PP1 inhibition from metaphase I onwards impairs meiosis I exit, but has no impact on meiotic abnormalities. (A) Schematic representation of the experimental design. Prophase I oocytes were released from meiotic arrest into standard culture medium for 8 h before being cultured in medium containing TMC (0.5-5 μM) or vehicle control (0.1% DMSO). (B) Graphical representation of meiotic stage based on phase microscopy at 16 h into culture. Bar graph shows the percentage of oocytes at each meiotic stage. n=50-52 oocytes over two replicates. * denotes a significant difference from Veh control oocytes (Fisher’s exact test). (C) Graphical representation of the meiotic stage and DNA morphology of oocytes at 16 h into culture. Solid colors represent normal DNA morphology for the meiotic stage, and hatched colors represent abnormal DNA morphology for the meiotic stage. n=33-40 oocytes over two replicates.

These surprising results prompted us to hypothesize that loss of PP1 activity specifically during the G2/M transition would have effects later in meiosis, such as on meiotic exit or metaphase II oocyte quality. To test this, oocytes were cultured in TMC during the G2/M transition (0 - 3 h post-meiotic arrest release), then were transferred into medium lacking TMC for the remainder of IVM (3 - 16 h post-meiotic arrest release). However, no difference in meiotic progression was observed in these oocytes, based on analyses of the ability of oocytes to progress through meiosis by phase microscopy and of DNA in DAPI-stained oocytes by fluorescence microscopy (Supp Fig. S3).

## Discussion

Temporal fluctuations in protein phosphorylations are essential for M-Phase entry and progression. Kinases have a well-established role in this process, however, the contribution and regulation of phosphatases during M-Phase is poorly understood. Here we establish that loss of oscillations in PP1 activity causes a range of severe meiotic defects, pointing to crucial roles for PP1 in oocyte meiosis and, more broadly, M-Phase regulation.

### PP1 and chromatin architecture

PP1 plays a key role in regulating chromatin architecture in a variety of systems (Bui et al., 2004; Gil and Vagnarelli, 2019). We found that PP1 inhibition and activation altered chromatin configuration at prophase I and metaphase I, respectively, suggesting that PP1 has important functions in chromatin architecture during oocyte meiosis. Prophase I oocytes have two chromatin configuration states: uncondensed euchromatin (NSN) or a condensed heterochromatin ring that surrounds the nucleolus (SN) (Bogolyubova and Bogolyubov, 2020). PP1 inhibition increased the proportion of prophase I oocytes with NSN chromatin configuration. As oocytes gain the ability to undergo the G2/M transition and exit from prophase I arrest (acquiring an ability known as meiotic competence), there is a shift from NSN to SN chromatin (Bellone et al., 2009; Zuccotti et al., 2002), with this shift accompanied by a translocation of PP1 to the nucleus (Smith et al., 1998; Wang et al., 2004). Post-translational modifications, including phosphorylation of H3, play a major role in heterochromatin formation. In somatic cells, dephosphorylation of S28 on H3 (H3S28) via the PP1 holoenzyme, PP1:RepoMan, is fundamental for heterochromatin formation (de Castro et al., 2017). H3S28 is phosphorylated in mouse prophase I oocytes, with pH3S28 dephosphorylated in a PP1/PP2A dependent manner at prophase I (Swain et al., 2007). Our findings, taken with these studies, suggest that PP1 plays a role in chromatin configuration changes at the NSN-to-SN transition, potentially via PP1-mediated H3S28 dephosphorylation. Therefore, it is possible that the observed change in chromatin configuration with PP1 inhibition is a product of SN heterochromatin decondensing, however, whether this results in a subsequent loss of meiotic competence is unknown.

Condensins and phosphorylation of H3 play a vital role in metaphase chromosome architecture (Fischle et al., 2005; Hirota et al., 2004; Hirota et al., 2005; Piskadlo et al., 2017). Our data show that PP1 activation reduced M-Phase phosphorylation of two PP1 substrates, H3 at T3 and at S10. During M-Phase, H3 is phosphorylated at T3 by Haspin and at S10 by Aurora B and C (Balboula and Schindler, 2014; Nguyen et al., 2014; Qian et al., 2011; Wang et al., 2016). Phosphorylation of H3 at T3 and S10 appears to play an important role in the recruitment of condensin to chromosomes. In *Drosophila*, Aurora kinases recruit condensin to chromosomes in mitosis, and in mouse oocytes active Haspin is required for Aurora C and condensin chromosome association at metaphase I (Giet and Glover, 2001; Nguyen et al., 2014). Notably, the abnormal pointy chromosome architecture observed with PP1 activation is phenocopied in mouse oocytes with oocyte-specific conditional loss of condensin and Haspin inhibition (Lee et al., 2011; Nguyen et al., 2014). Therefore, activation of PP1, and subsequent reduction of pH3T3 and pH3S10 may lead to loss of chromosome condensin, ultimately causing the observed chromosome abnormalities. Further studies are warranted to determine how PP1 regulates chromosome architecture during M-Phase.

### Active pools of PP1 in prometaphase impact M-Phase exit and oocyte quality

In mitosis, PP1 plays a pivotal role in the metaphase/anaphase transition (e.g., silencing the spindle assembly checkpoint and APC/C^CDC20^ activation) and cytokinesis. In human mitotic cells, PP1 inhibition or siRNA-mediated loss of PP1 leads to delayed metaphase/anaphase transition, metaphase arrest, and binucleated cells (a sign of cytokinesis failure) (Bancroft et al., 2020; Bhowmick et al., 2019; Capalbo et al., 2019; Cheng et al., 2000; Winkler et al., 2015). We found a similar phenotype in TMC-treated oocytes, with PP1 inhibition for the duration of meiosis (0-16 h post-meiotic arrest release) causing metaphase I arrest or meiotic abnormalities (e.g., chromosome misalignment at metaphase II or cytokinetic failure (two spindles without a PB)). On the other hand, TMC-treatment from metaphase I-to-metaphase II arrest (8-16 h post-meiotic arrest release) showed remarkably few abnormalities, with oocytes largely able to form a normal metaphase II oocyte by 16 h. These disparate results point to an exciting new role for PP1 during prometaphase that establishes the ability of oocytes to undergo metaphase I/anaphase I transition and form a normal metaphase II oocyte. Approximately 40% of PP1 escapes the CDK1-mediated inhibitory phosphorylation of T320, such that there is a substantial subpopulation of PP1 that could be active to mediate this novel function for PP1 in prometaphase (Olsen et al., 2010). Our data further confirm the presence of an active pool of PP1, with a significant increase in pH3T3 observed in oocytes treated with TMC during prometaphase (0-7 h post-meiotic arrest release).

There are two probable causes of the abnormalities observed in oocytes treated with TMC for the duration of meiosis (0-16 h): (1) continued phosphorylation of prometaphase PP1 substrates, and (2) altered protein stability. Prometaphase PP1 substrates have been revealed in a recent study of meiotic phosphorylation dynamics in starfish oocytes, with a subset of CalA (PP1/PP2A inhibitor) sensitive protein phosphorylations peaking at NEBD and decreasing by metaphase I (Swartz et al., 2021). In oocytes with PP1 inhibition for the duration of meiosis (0-16 h), these early M-Phase PP1 phospho-sites could be maintained, likely contributing to the defects observed in these oocytes. In contrast, in oocytes with PP1 inhibition only during metaphase I (8-16 h), these early M-phase PP1 phospho-sites would not be affected, thus being a possible explanation for the differences in the effects of TMC treatments during these two time windows. Protein phosphorylation, including phosphorylation affected by PP1, also plays a role in proteasomal protein targeting, and therefore protein half-life. PP1-mediated dephosphorylation prevents proteasome-mediated degradation of specific proteins (PER2 and MYC) (Dingar et al., 2018; Gallego et al., 2006). Additionally, PP1-mediated dephosphorylation of CDC20 plays a role in activating APC/C^CDC20^, the major E3 ligase involved in the degradation of pro-M-Phase proteins at metaphase/anaphase transition (Bancroft et al., 2020; Kim et al., 2017). Degradation of the APC/C^CDC20^ substrates securin and cyclin B starts in prometaphase of oocytes and is essential for metaphase I/anaphase I transition and accurate chromosome segregation (Levasseur et al., 2019; Thomas et al., 2021). Consequently, PP1 inhibition in oocytes at prometaphase may alter the degradation kinetics of proteins, impacting M-Phase progression. Our results show that an active pool of PP1 during prometaphase is essential for M-Phase exit and the formation of normal metaphase II oocytes. The specific PP1 holoenzymes and their substrates involved in these prometaphase events remain to be identified.

### PP1 activity is needed to maintain metaphase I chromosome congression

PP1 inhibition during prometaphase (0-7 h post-meiotic arrest release) appears to accelerate chromosome alignment compared to controls. However, this chromosome alignment is lost in oocytes that fail to exit metaphase I by 16 h of maturation. A possible explanation for this observation relates to PP1’s kinetochore functions. At the kinetochores, PP1 is in a feedback loop with PP2A-B56. The changing activity of these phosphatases has essential roles in chromosome alignment and microtubule-kinetochore attachments (Vallardi et al., 2017). PP2A-B56 initializes microtubule-kinetochore attachments, aligns chromosomes, and recruits PP1 to the kinetochore, with PP1 then stabilizing microtubule-kinetochore attachments and removing PP2A-B56 from the kinetochore (Liu et al., 2010; Nijenhuis et al., 2014; Xu et al., 2013). Thus, PP1 inhibition may favor kinetochore PP2A-B56 and accelerate chromosome congression initially (as observed at 7 h), but then later, failure to stabilize microtubule-kinetochore attachments could cause chromosome misalignment in metaphase I oocytes at 16 h.

### Timing of PP1 activation impacts oocyte survival

As discussed above, the timing of PP1 inhibition significantly impacts M-Phase outcomes in oocytes. Similarly, the sensitivity of oocytes to PP1 activation is dependent on M-Phase stage. PP1 activation at G2/M transition (0-5 h post-meiotic arrest release) reduces oocyte viability significantly more than PP1 activation during M-Phase (3-8 h post-meiotic arrest release). Prophase I arrest (equivalent to G2 in mitosis) is characterized by high phosphatase activity and low kinase activity, particularly kinases that drive M-Phase (e.g., CDK1) (Swartz et al., 2021). Conversely, high kinase activity and low phosphatase activity maintains M-Phase progression. CDK1 is key to establishing this high kinase / low phosphatase environment, with CDK1-mediated phosphorylations activating other pro-M-Phase kinases and inhibiting phosphatases (Crncec and Hochegger, 2019; Moura and Conde, 2019; Rata et al., 2018). PP1 is part of this kinase/phosphatase interplay; at M-Phase entry, CDK1 phosphorylates PP1 at T320, which inhibits PP1 activity (Dohadwala et al., 1994; Kwon et al., 1997). Conversely, auto-dephosphorylation of T320 reactivates PP1 (Wu et al., 2009). Therefore, it is probable that PP1 activation during M-Phase is less deleterious as compared to the G2/M transition because in M-phase, (a) PP1 has to remove the CDK1-mediated inhibitory phosphorylation of T320, and (b) PP1 has to counter the activity of kinases that have already established high activity. In contrast, at the G2/M transition, high PP1 activity is expected to prevent activation of M-Phase kinases leading to an environment of significantly higher phosphatase activity and lower kinase activity than observed in PDP-Nal-treated oocytes at M-Phase. In support of this hypothesis, PDP-Nal treatment caused a 2-to-2.5-fold greater loss of PP1 substrates pH3T3 and pH3S10 at G2/M transition (0-5 h) compared to M-Phase (3-8 h).

## Conclusion

In summary, we found that (a) PP1 has critical roles at multiple stages of mammalian oocyte meiosis, and (b) impairing the anticipated oscillations in PP1 activity negatively impacts M-Phase. Importantly, we show for the first time that the timing of PP1 inhibition/activation significantly influences meiotic outcomes. Our study also highlights the advantage of mouse oocytes for studying PP1 at discrete subphases of M-Phase (e.g., prometaphase vs. metaphase). The extended time frame of M-Phase in mouse oocytes (~10 h from G2/M to M-Phase exit) allows for the investigation of PP1’s roles at specific subphases of M-Phase that is not currently possible in M-Phase of mammalian cell lines. Demonstrating the importance of this evaluation of PP1 at specific subphases of M-Phase, our study found a novel role for PP1 in prometaphase that is essential for M-Phase exit. Taken together, our results establish the vital role PP1 plays in mammalian oocyte meiosis, and therefore, female fertility. More broadly, this study underscores the need for accurate phosphatase regulation during M-Phase.

## Methods

### Animals and Ethics Approval

6-11-week-old female CF-1 mice (Envigo) were used in accordance with Purdue Animal Care and Use Committee guidelines. Mice were housed under a 12 h light: dark cycle with ad libitum water and food.

### Oocyte Collection and *In vitro* maturation

Prophase I stage oocytes were collected into Whitten’s-HEPES medium (109.5 mM NaCl, 4.7 mM KCl, 1.2 mM KH_2_PO_4_, 1.2 mM MgSO_4_, 5.5 mM glucose, 0.23 mM pyruvic acid, 4.8 mM lactic acid hemicalcium salt, with 7 mM NaHCO_3_, 15 mM HEPES) supplemented with 0.25 mM dbcAMP (Sigma-Aldrich, D0627) or 2.5 μM milrinone (Sigma-Aldrich, M4659) to maintain prophase I arrest. Cumulus cells were mechanically removed via repeated pipetting of oocytes before transfer into Whitten’s bicarbonate (Whitten’s medium without HEPES and with 22 mM NaHCO3) supplemented with dbcAMP or milrinone. For each experiment, oocytes were pooled from 3-7 females.

For *in vitro* maturation (IVM), oocytes were washed into Whitten’s-Bicarb without dbcAMP or milrinone to release oocytes from prophase I arrest. Oocytes were matured for up to 16 h at 37°C, 5% CO_2_. For assays to evaluate inappropriate exit from prophase I arrest, oocytes were washed into Whitten’s-Bicarb supplemented with 0.2 μM milrinone for 16 h at 37°C, 5% CO_2_.

### Small molecule inhibitors and activators

Oocytes were cultured in tautomycetin 0.5 - 5 μM (TMC; selective PP1c inhibitor; Bio-Techne, 2305), 2.5 - 40 μM PDP-Nal (PP1c activator; a gift from Maja Köhn), 50 nM Calyculin A (CalA; Cell Signaling Technology (CST), 9902S) or vehicle (0.1% DMSO, equal to DMSO in 5 μM TMC treatment group) for the duration of IVM or for discrete windows of time (see Fig. 1C). For experiments that added TMC or PDP-Nal during meiosis I (*e.g.*, 3 h post-meiotic arrest release), oocytes still at prophase I arrest at 2.5 h post-dbcAMP/milrinone washout were removed. Finally, for oocytes with TMC treatment for 7 and 9 h post-meiotic arrest release, oocytes still at prophase I arrest were removed 6.5 h post dbcAMP/milrinone washout. Inhibitor/activator concentration and culture conditions for individual experiments are detailed in the results section and figure legends.

### Immunofluorescence and F-actin staining

Oocytes were fixed in 4% paraformaldehyde (Sigma-Aldrich, P6148) in HEPES buffer (130 mM KCl, 25 mM HEPES, 3 mM MgCl_2_, and 0.06% Triton-X, pH 7.4) for 30 minutes at 37°C prior to permeabilization in PBS with 0.1% Triton X-100 (Fisher Scientific, BP151–500) and blocking in PBS with 0.1% BSA (Sigma-Aldrich, A9647) and 0.01% Tween-20 (Sigma-Aldrich, P7949) at room temperature. Oocytes were incubated with primary antibodies for α-tubulin (Zymed, clone Z023, 3605884, 1:100, or Developmental Studies Hybridoma Bank, clone 12G10, 10.6 μg/ml), pH3T3 (CST, 13576S, 220 ng/ml), pH3S10 (CST, 9701S, 25 ng/ml), or pAKT(S473) (CST, 9271T, 200 ng/ml) for 16 h at 4°C. After 16 h, oocytes were washed in PBS with 0.1% BSA and 0.1% Tween-20 and incubated with 7.5 μg/ml goat anti-mouse IgG or donkey anti-rabbit conjugated to Alexa Fluor 488 (Jackson ImmunoResearch) at room temperature. For pAKT(S473) immunofluorescence 100 μM Na_3_VO_4_ (Sigma-Aldrich, S6508) was included in all buffers. For F-actin staining, 140 nM Acti-stain 555 phalloidin (Cytoskeleton Inc., PHDH1-A) was added to the secondary antibody. Oocytes were mounted in VectaShield containing 0.75 μg/ml 4-6-diamidino-2-phenylindole (DAPI; Sigma-Aldrich, D9542). Imaging was performed using a Zeiss Axio Observer Z1 microscope with AxioCam MRm Rev3 camera and ApoTome optical sectioning (Carl Zeiss, Inc.). Optical sectioning of oocytes was performed to include all DNA and associated microtubules within the cytoplasm with an optical distance of 1 μm between sections.

### Immunofluorescence analysis

Image analysis was performed with ImageJ Fiji distribution (Freeware; National Institutes of Health - https://imagej.nih.gov/ij/). Details for each type of analysis are specified below.

Chromatin configuration and chromosome alignment: Slides were blinded, and oocytes were imaged and assessed for DNA morphology and chromosome alignment to classify stage of meiosis (*e.g.,* prophase I, metaphase I, metaphase II) and chromosome alignment on the metaphase plate. Oocytes were classified as aligned if all DNA congressed on the metaphase plate, as mild chromosome misalignment if one to two chromosomes had not congressed to the metaphase plate, as moderate misalignment if three to four chromosomes had not congressed to the metaphase plate, or as severe misalignment if >4 chromosomes had not congressed to the metaphase plate or no metaphase plate was present.

Histone H3 phosphorylation fluorescence intensity (pH3T3 and pH3S10): DNA and histone H3 immunofluorescence optical sections were merged into a single DNA or histone H3 immunofluorescence Z-projection. Image threshold and analyze particles Fiji functions were used to determine DNA region-of-interest on DNA Z-projections automatically. Calculations of the pH3T3 and pH3S10 fluorescence intensity were then performed by first obtaining the value for the background signal (by multiplying the DNA area by the average of the mean background signal), and then subtracting this background signal from the integrated density value of the phospho-Histone H3 signal. Control oocytes were then normalized to an average value of 1, with this used to normalize all other groups. For experiments establishing H3T3 phospho-dynamics through meiosis (Fig. 6A), the fluorescence intensity of pH3T3 was performed on all oocytes regardless of their meiotic stage. For all other experiments quantifying pH3T3 and pH3S10 dynamics, only the fluorescence intensity of prometaphase or metaphase I oocytes were measured.

AKT(S473) phosphorylation fluorescence intensity: The integrated density of a single optical section of the whole oocyte pAKT(S473) immunofluorescence was measured. Fluorescence intensity was then calculated as described above and normalized to DMSO at 3 h to an average value of 1.

Distance of metaphase I plate to the cortex: A Z-projection of the DNA optical sections and the central F-actin optical section were merged into a single image. The metaphase I spindle migrates to the cortex in a pole-first manner (Supp Fig. 2B) (Fabritius et al., 2011). Therefore, a straight line from the center of the metaphase I plate to the closest region of the cortex with this pole-first directionality was measured (Supp Fig. 2C).

### Statistical Analysis

GraphPad Prism 9.2.0 was used for statistical analysis (GraphPad Software, www.graphpad.com), with a p-value of <0.05 considered statistically significant. For categorical data, the Fisher’s exact test was used. For continuous/numerical data, D’Agnostino-Pearson omnibus normality test was performed to determine if the data followed a normal distribution. For normally distributed numerical data, ANOVA with Tukey’s post hoc test was used. For all other data, the Kruskal-Wallis test with Dunn’s post hoc was used. The exact p-values for each analysis can be found in the results section. Additionally, further information on statistical tests used for each dataset can be found within the figure legends.

## Acknowledgments

This work was supported by funds from NIH (grant HD087561) and Purdue University to JPE. The authors gratefully acknowledge Malgorzata Trebacz and Maja Köhn (University of Freiburg) for providing the PP1 activators and control (PDP-Nal and PDPm-Nal). We also thank Brittany Allen-Petersen and Jennifer Ikle for critical reading of the manuscript.

---

**Supp Figure 1: PP1 activation from prometaphase to metaphase of meiosis I does not impact metaphase I plate position or chromosome alignment.** (A) Schematic representation of the experimental design. Prophase I oocytes were released from meiotic arrest and cultured for 3 h before treatment with increasing concentrations of PDP-Nal (2.5-10 μM), PDPm-Nal (10 μM) or standard culture medium (Ctrl; 0 μM PDP-Nal). (B) Schematic representation of the metaphase I spindle migration during meiosis I. Purple dashed arrow denotes pole-first directionality of the spindle migration. (C) Representative image of metaphase I plate to cortex measurement. Orange dashed line denotes distance measured, scale = 10 μm. (D) Graphical comparison of metaphase plate distance to oocyte cortex (Kruskal-Wallis test with Dunn’s post hoc). n=34-60 oocytes over 4-5 replicates. Scatter dot plots show each individual oocytes metaphase plate distance to cortex plus mean and SEM. (E) Graphical representation of chromosome alignment in metaphase I oocytes (Fisher’s exact test). n=43-69 oocytes over 4-5 replicates.

**Supp Figure 2: pH3T3 expression at specific stages of meiotic maturation.** Representative images of oocyte pH3T3 fluorescence, DNA and DIC at metaphase I (Met I; 7-8 h), anaphase I/telophase I (ana I/telo I; 9-10 h), cytokinesis, and early metaphase II (met II forming; 10h). Time denotes the number of hours into meiotic maturation that individual stages of meiosis were observed. Orange dashed line denotes insert region, scale = 10 μm.

**Supp Figure 3: PP1 inhibition during meiosis I does not impact metaphase I plate position.** (A) Schematic representation of the experimental design. Prophase I oocytes were released from meiotic arrest and cultured in medium containing TMC (5 μM) or vehicle control (0.1% DMSO) for 7 or 9 h. (B) Graphical comparison of metaphase I plate distance to oocyte cortex at 7 or 9 h post-meiotic arrest release (Kruskal-Wallis test with Dunn’s post hoc). n=27-55 oocytes over two replicates. Scatter dot plots show each individual oocytes metaphase plate I distance to cortex plus mean and SEM.

**Supp Figure 4: PP1 inhibition at G2/M transition only does not cause significant meiotic abnormalities.** (A) Schematic representation of the experimental design. Prophase I oocytes were released from meiotic arrest and cultured for 3 h in medium containing TMC (0.5-5 μM) or vehicle control (0.1% DMSO) before being washed into standard culture medium (no TMC or DMSO). (B) Graphical representation of meiotic stage based on phase microscopy at 16 h into culture. Bar graph shows the percentage of oocytes at each meiotic stage. n=47-49 oocytes over two replicates. (C) Graphical representation of the meiotic stage and DNA morphology of DAPI-stained oocytes at 16 h into culture. See Figure 2 for examples and descriptions. Solid colors represent normal DNA morphology for the meiotic stage, and hatched colors represent abnormal DNA morphology for the meiotic stage. n=27-36 oocytes over two replicates.

## Notes

### Competing Interest Statement

The authors have declared no competing interest.

### Summary of Updates

Introduction updated to shorten and clarify; Results section "Tautomycetin (TMC) selectively inhibits PP1 in mouse oocytes" and accompanying figure added; Rationale for quantification of chromosome alignment and spindle to cortex distance added; Supplemental figures updated.

