## Supplementary figures and images for "M-Phase oscillations in PP1 activity are essential for accurate progression through mammalian oocyte meiosis"

### Supp Fig 1

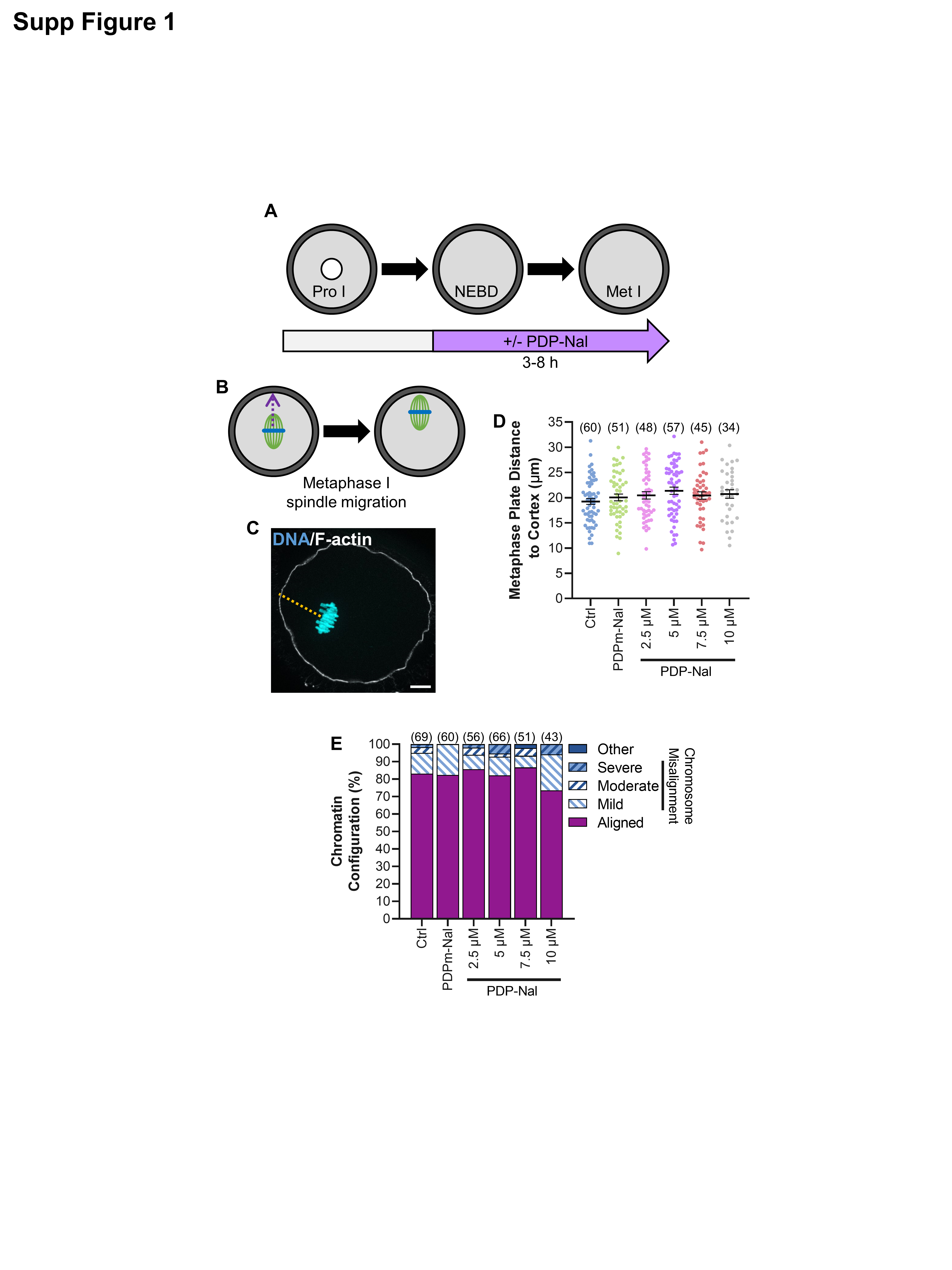

### Supp Fig 2

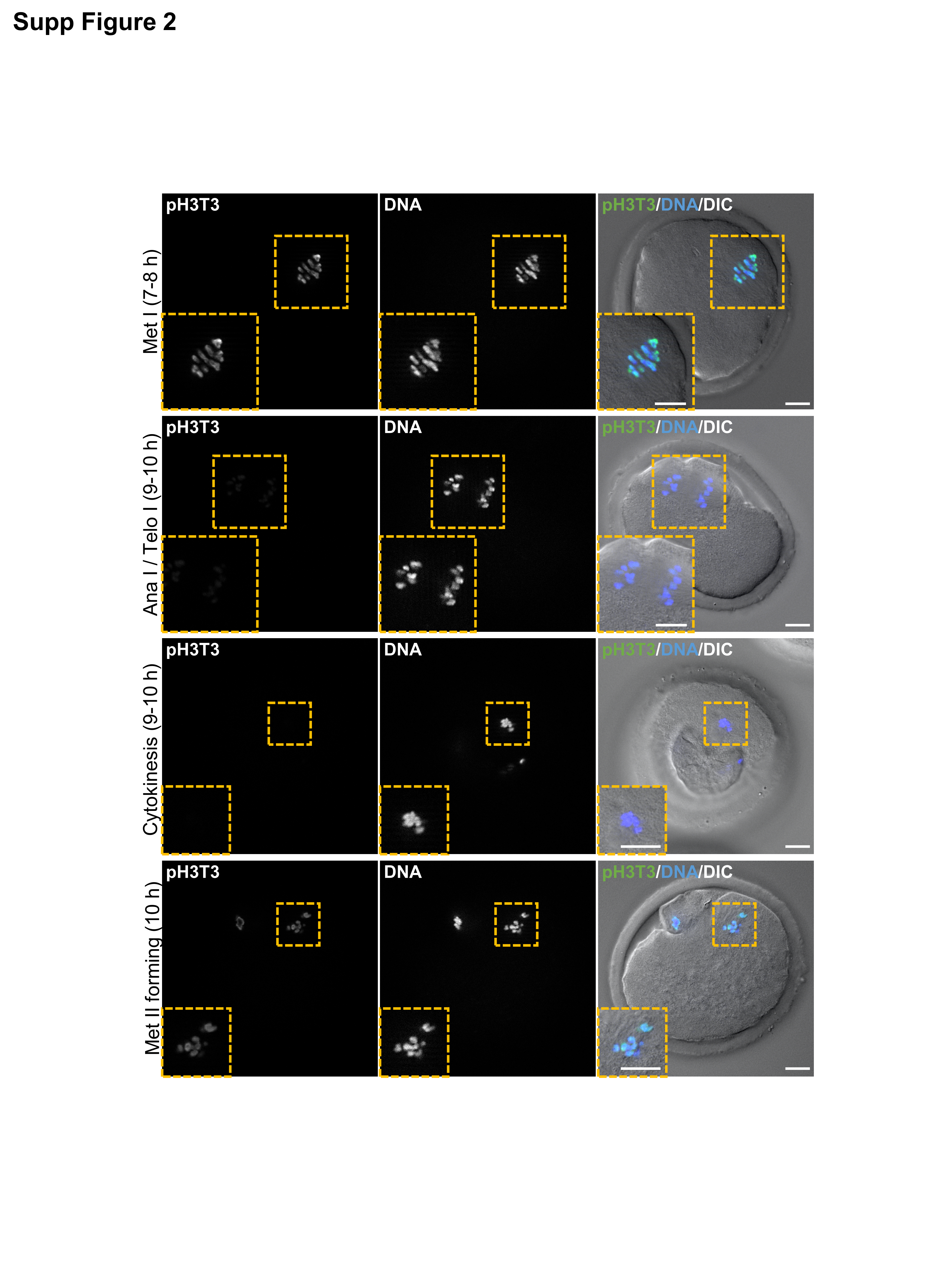

### Supp Fig 3

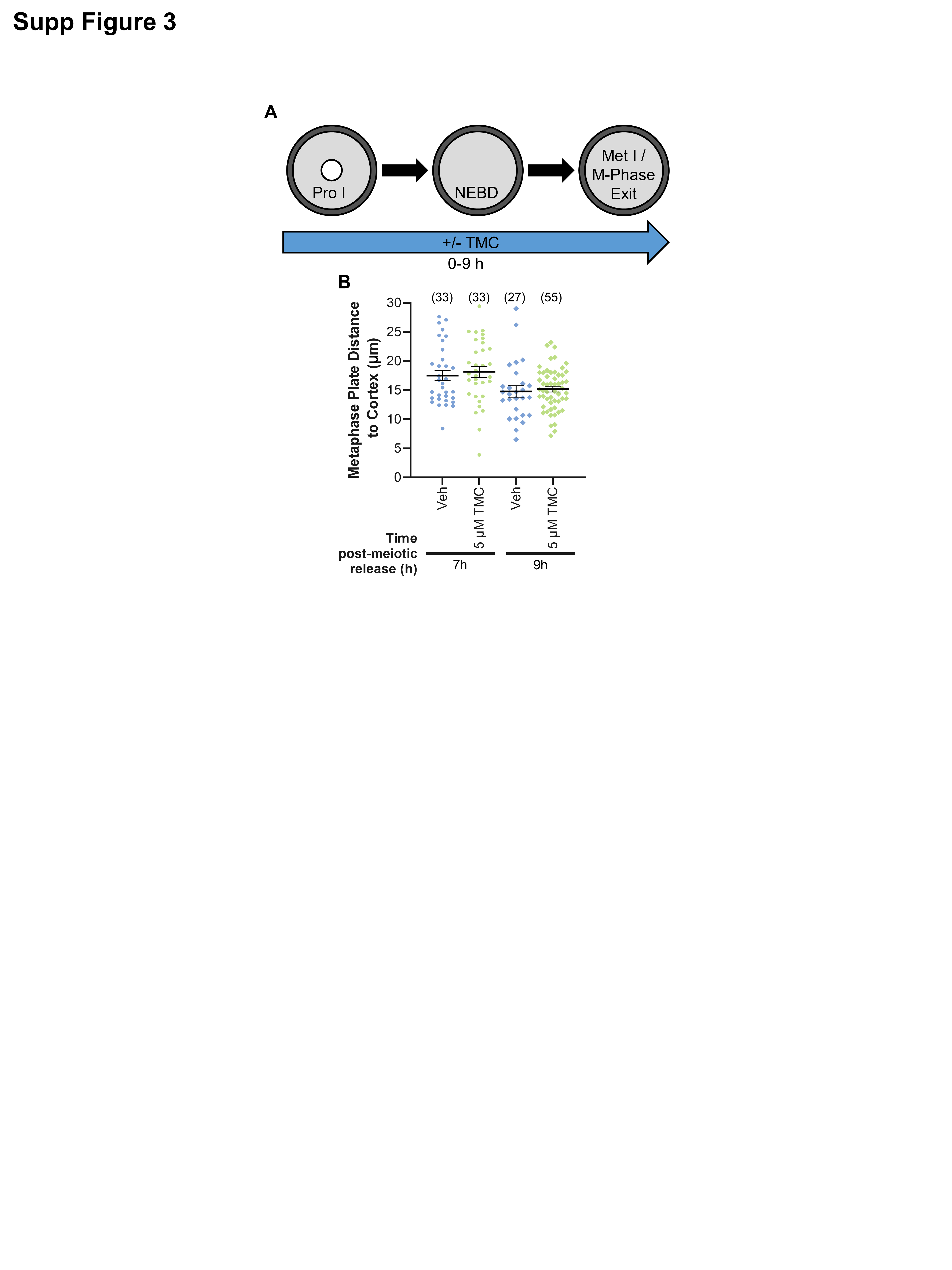

### Supp Fig 4

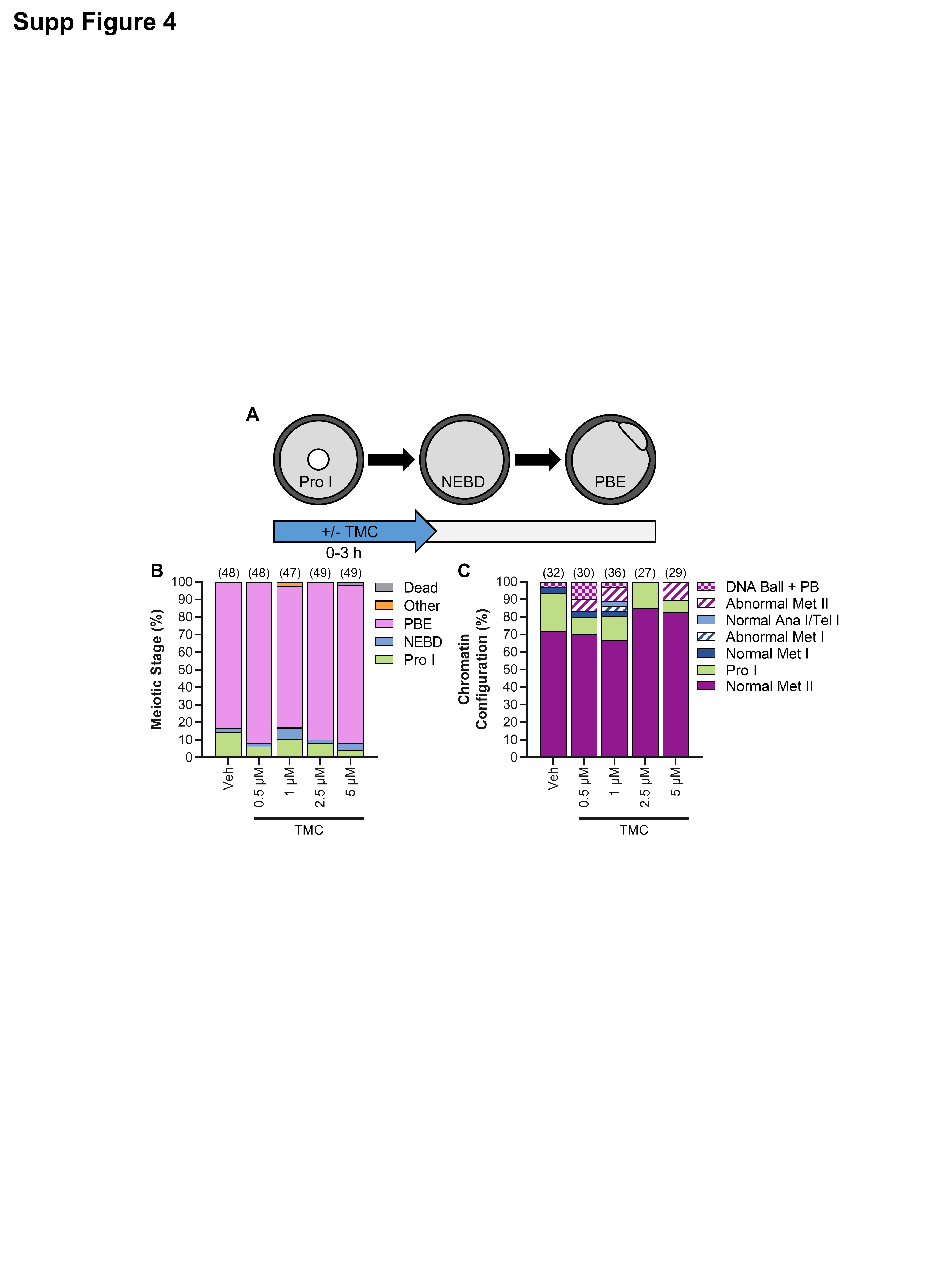
